# Immunofilaments Provide a Nanoscale Platform for In Vivo T Cell Expansion and Cancer Immunotherapy

**DOI:** 10.1101/2022.10.14.512109

**Authors:** Lea Weiss, Jorieke Weiden, Yusuf Dölen, Emilia M. Grad, Eric A.W. van Dinther, Marjolein Schluck, Loek J. Eggermont, Guido van Mierlo, Uzi Gileadi, Ariadna Bartoló-Ibars, René Raavé, Mark A.J. Gorris, Kiek Verrijp, Michael Valente, Bart Deplancke, Martijn Verdoes, Daniel Benitez-Ribas, Vincenzo Cerundolo, Sandra Heskamp, Annemiek B. van Spriel, Carl G. Figdor, Roel Hammink

## Abstract

Adoptive T cell therapy has successfully been implemented for the treatment of cancer. Nevertheless, the precarious ex vivo expansion of T cells by artificial antigen-presenting cells (aAPCs) remains cumbersome and can compromise T cell functionality, thereby limiting their therapeutic potential. Here, we propose a radically different approach aiming at direct expansion of T cells in vivo, thereby omitting the need for large-scale ex vivo T cell production. We engineered nanosized immunofilaments (IFs), consisting of a soluble semiflexible polyisocyanopeptide polymer backbone that presents peptide-loaded major histocompatibility complexes and co-stimulatory molecules in a multivalent fashion. We demonstrate that IFs readily activate and expand antigen-specific T cells in a manner highly similar to natural APCs, as evidenced by transcriptomic analyses of T cells. Upon intravenous injection, IFs reach lymphoid organs including spleen and lymph nodes and induce antigen-specific T cell responses in vivo. Moreover, IFs display remarkable anti-tumor efficacy resulting in inhibition of melanoma metastases formation and reduction of primary tumor growth in synergy with immune checkpoint blockade. In conclusion, nanosized IFs represent a powerful new type of aAPC that provide a modular platform for direct activation and expansion of antigen-specific T cells in vivo, which can greatly contribute to cancer immunotherapy.

## 1. Introduction

To date, the cancer immunotherapy field is dominated by therapeutic strategies that aim to exploit the cytotoxic potential of T cells. T cells can trigger tumor cell death following recognition of tumor antigens presented as peptide epitopes on major histocompatibility complexes (pMHC) on the cell surface. In particular, unleashing existing tumor-directed T cell responses by administrating monoclonal antibodies that block co-inhibitory receptors PD-1 and/or CTLA-4 on T cells has proven highly beneficial, and is termed immune checkpoint blockade.^[1–4]^ To further enhance the proportion of cancer patients that respond to immunotherapy, these strategies are complemented with treatment modalities aimed at expanding the number of already existing T cells or by inducing novel tumor-specific T cells. To this end, cancer patients are treated with adoptive T cell therapy (ACT), in which they receive infusions of ex vivo-expanded tumor-infiltrating lymphocytes (TILs) or genetically engineered chimeric antigen receptor (CAR)-expressing T cells.

Artificial antigen-presenting cells (aAPCs) provide an essential off-the-shelf tool for the ex vivo expansion of T cells for ACT. These aAPCs mimic the interaction between natural APCs such as dendritic cells (DCs) and T cells during T cell priming. To this end, synthetic constructs have been designed that present molecular signal to T cells in a controlled manner to 1) trigger T cell receptor (TCR) signaling by presenting agonistic anti (α)CD3 antibodies or antigen-specific pMHC, and 2) provide secondary signals by triggering co-stimulatory receptors such as CD28 to enhance and prolong T cell responses. In addition, cytokines such as interleukin-2 (IL-2) or type I interferons can be provided as a third signal to promote T cell survival, control T cell phenotype and enhance T cell functionality. The most notable example of aAPCs that are applied in the clinical settings are micrometer-sized iron oxide particles (Dynabeads), which typically present αCD3/αCD28 antibodies to foster ex vivo polyclonal T cell expansion. Various alternative synthetic systems have been developed that support more rapid T cell expansion or promote a favorable T cell functionality or phenotype.^[5–9]^ However, large-scale ex vivo multiplication of T cells remains laborious and costly, and can also compromise T cell functionality and viability, leading to a suboptimal therapeutic efficacy for current ACT strategies.^[10–13]^ Here, we propose a radically different approach by designing nanosized soluble aAPCs that facilitate systemic administration and that can expand (adoptively transferred) tumor-specific T cells directly in vivo, thereby omitting the need for ex vivo T cell production. We previously demonstrated that nanosized immunofilaments (IFs) effectively stimulate polyclonal T cell responses ex vivo. These soluble IFs are based on filamentous synthetic polyisocyanopeptides (PIC) which are rather long (∼400 nm) but very thin (∼1-2 nm) polymers and are semiflexible.^[14,15]^ Owing to their length, IFs can be functionalized with multiple biomolecules such as antibodies and cytokines through azides that are incorporated into the polymer side chains. Immunofilaments decorated with T cell-stimulating antibodies αCD3/αCD28 or αCD3 combined with immobilized cytokines were found to induce strong T cell activation and expansion ex vivo.^[16,17]^ Co-presentation of αCD3 and αCD28 on the same polymer furthermore outperformed single antibodies presented on separate polymers, suggesting that combined multivalent presentation of T cell-stimulating signals is critical. In addition, we observed that these semiflexible IFs induced T cell responses were superior compared to those evoked by more rigid substrates.^[16,18]^ This underlines the importance of molecular flexibility of nanosized systems to facilitate the presentation of biomolecular signals to T cells in a multivalent fashion to induce potent T cell responses.^[16,18]^

So far, these IFs have only been applied for polyclonal T cell stimulation. Here, we demonstrate that nanosized IFs functionalized with pMHC and co-stimulatory molecules (αCD28 or IL-2) (**Figure 1**) induce strong antigen-specific T cell activation, both ex vivo and in vivo. Furthermore, IFs accumulated in lymphoid organs upon intravenous injection and displayed remarkable antitumor efficacy in vivo, as they inhibited metastases formation and reduced primary tumor growth in combination with immune checkpoint blockade. We conclude that IFs constitute a modular platform that can be applied as a unique nanosized tool to trigger robust anti-cancer immune responses in vivo.

**Figure 1.**
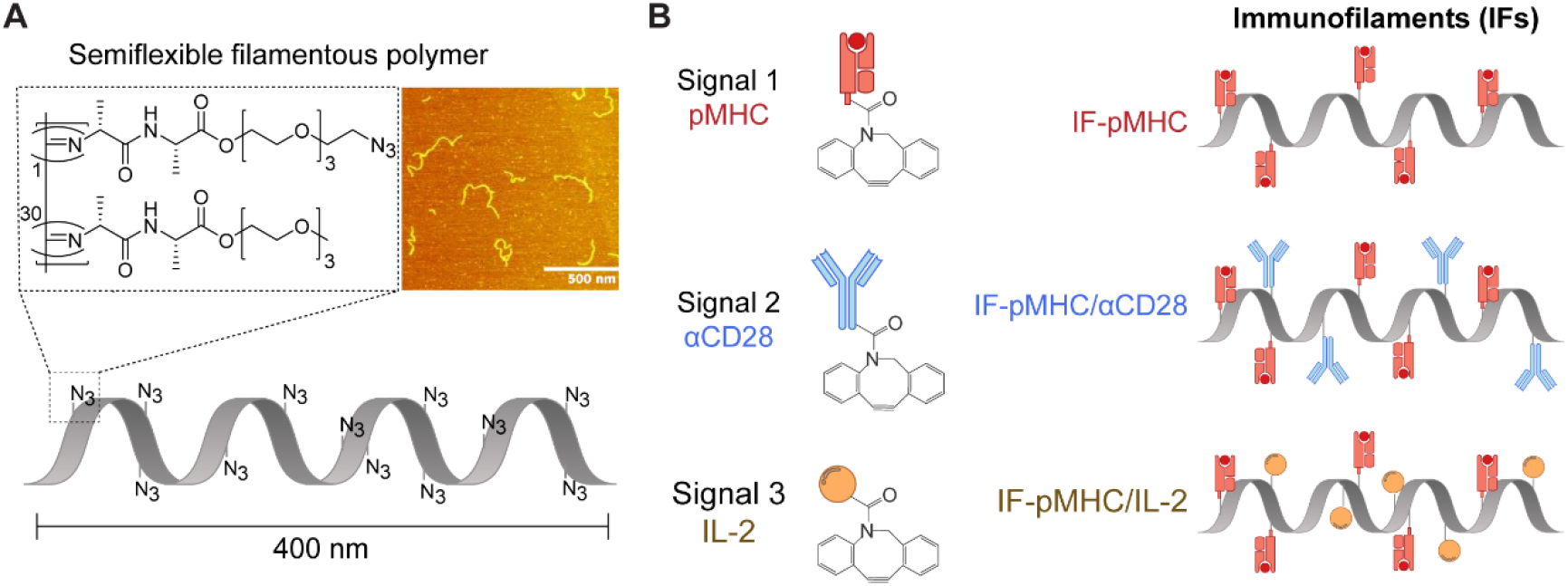
Schematic overview of the preparation of nanosized immunofilaments (IFs). (A) Chemical structure of azide-functionalized polyisocyanopeptides (PIC) (left) and atomic force microscopy image of PIC (right) with an average length of ∼400 nm and thickness of ∼2nm. (B) Different types of IFs are prepared by conjugating dibenzocyclooctyne-functionalized biomolecules (pMHC, αCD28, IL-2) to azide-functionalized PIC.

## 2. Results and Discussion

### 2.1 Design of PIC-based IFs

Immunofilaments for antigen-specific T cell stimulation were prepared by conjugating T cell-activating biomolecules to nanosized PIC polymers. First, azide-functionalized PIC were synthetized by co-polymerization of azide-terminated isocyanopeptide monomers and non-functional methoxy-terminated monomers (1/30 ratio) as described previously.^[17,19,20]^ We obtained semiflexible PIC polymers that were ∼400 nm in length and 2 nm in width as measured by atomic force microscopy (Figure 1, inset) with a previously established statistical average of one azide group every 3.5 nm.^[16,18,21]^ Half of the azides were converted into biotin to enable affinity-based purification of the PIC polymers.^[22]^ The remaining azides were used to attach dibenzocyclooctyne (DBCO)-modified T cell-activating proteins via strain-promoted azide-alkyne cycloaddition.^[23]^ Using this strategy, we prepared IFs with pMHC alone or combined with agonistic anti-mouse CD28 or recombinant IL-2 (Figure 1). The complete characterization and specifications of these polymers are provided in **Table 1** and **Table 2**.

**Table 1.** Concentration of IFs and protein spacing used for ex vivo experiments.

| Immunofilament | Concentration of signal 1 | Protein spacing on IFs |
| --- | --- | --- |
| IF-pMHC<sup>(SIIN)</sup> | 2.5 ng/mL | 54 nm |
| IF-pMHC<sup>(SIIN)</sup>/αCD28 |  | 43 nm/ 140 nm |
| IF-pMHC<sup>(SIIN)</sup>/IL-2 |  | 43 nm/ 72 nm |
| IF-pMHC<sup>(SIIN)</sup>/αCD28/IL-2 |  | 49 nm/ 158 nm/ 79 nm |
| IF-pMHC<sup>(SIIT)</sup> | 25 ng/ml<br>(RNAseq: 5 ng/mL) | 177 nm |
| IF-pMHC<sup>(SIIT)</sup>/αCD28 |  | 137 nm/ 87 nm |
| IF-pMHC<sup>(SIIT)</sup>/IL-2 |  | 137 nm/ 127 nm<br>(RNAseq: 90 nm/ 64 nm) |
| IF-A2<sup>(NYESO)</sup> | 500 ng/mL (1G4 T cells) | 139 nm or 57 nm |
| IF-A2<sup>(NYESO)</sup>/IL-2 | 750 ng/mL (NY-ESO-1 TCR-transfected human T cells) | 124 nm/ 80 nm or 37 nm/ 29 nm |
| IF-ProL | 2 μg/mL | 39 nm |
| IF-ProL/αCD28 |  | 20 nm/ 49 nm |
| IF-ProL/αCD28/IL-2 |  | 59 nm/ 142 nm |

**Table 2.** Concentration of IFs and protein spacing used for in vivo experiments.

| Immunofilament | Amount of signal 1 (ng) | Protein spacing on IFs | Corresponding Figure |
| --- | --- | --- | --- |
| IF-pMHC<sup>(SIIN)</sup><br>or free pMHC | 290 | 69 nm | Figure 6A-D, Figure 7 B,C |
| IF-pMHC<sup>(SIIN)</sup> | 290 | 35 nm | Figure S6A-E, Figure S7 B,C |
| IF-pMHC<sup>(SIIN)</sup> | 100 – 1000 | 47 nm | Figure S7I,J |
| IF-pMHC<sup>(SIIN)</sup> LD | 1000 | 87 nm | Figure 7D,E |
| IF-pMHC<sup>(SIIN)</sup> HD | 1000 | 26 nm | Figure S7E-G |
| IF-pMHC<sup>(SIIN)</sup> | 100-1000 | 35 nm | Figure S7L,M |
| IF-pMHC<sup>(SIIN)</sup> | 1400 | 50 nm | Figure 8B,D |
| IF-pMHC<sup>(SIIN)</sup> | 500 | 58 nm | Figure 8F-I |
| IF-A2<sup>(NY-ESO-V)</sup>/IL-2 | 1400 | 43 nm / 58 nm | Figure S8 |

### 2.2 Antigen-specific activation of primary mouse OT-I T cells by IFs ex vivo

To validate that IFs can be used for antigen-specific stimulation, we used IFs presenting mouse H-2Kb-SIINFEKL (IF-pMHC^(SIIN)^) to activate primary OT-I CD8^+^ T cells, which express a transgenic TCR specific for the ovalbumin (OVA) epitope SIINFEKL (**Figure 2A**). The IF-pMHC^(SIIN)^ rapidly activated OT-I cells, resulting in expression of activation marker CD25 (**Figure 2B, Figure S1A**), production of IL-2 (**Figure S1B**) and interferon gamma (IFNγ) (**Figure 2C**) trending towards higher levels when compared with T cells that were exposed to free pMHC^(SIIN)^. The IF-pMHC^(SIIN)^ also induced strong T cell proliferation at levels similar to T cells stimulated with free pMHC^(SIIN)^ (**Figure S1C**,**D**) and acquired the ability to lyse OVA-expressing B16 melanoma cells with high efficiency (**Figure S1E**). These data demonstrate the potential of nanosized IFs to expand functional T cells.

**Figure 2.**
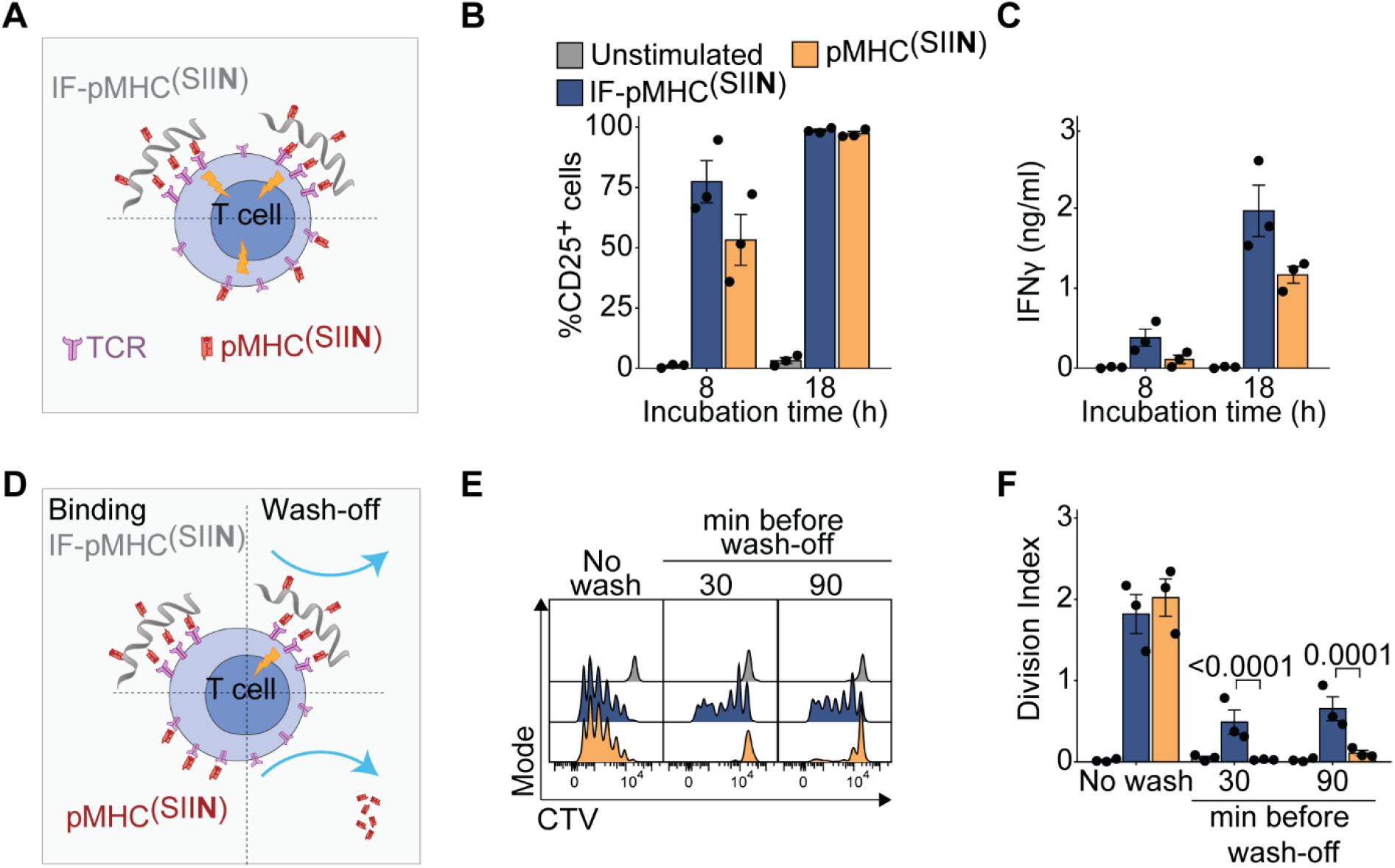
Multivalent IFs presenting pMHC activate and expand antigen-specific primary mouse T cells ex vivo. (A) Schematic overview of IF-pMHC^(SIIN)^ and free pMHC^(SIIN)^ for stimulation of murine OT-I T cells. (B) Flow cytometry quantification of the percentage of activated CD25^+^ OT-I T cells at different time points. Statistical significance was tested with two-way ANOVA on logit-transformed data. (C) The production of IFNγ by OT-I T cells at different time points was determined by ELISA. Statistical significance was tested with two-way ANOVA on log-transformed data. (D) Schematic overview of an experiment where IF-pMHC^(SIIN)^ and free pMHC^(SIIN)^ are washed off from OT-I T cells after 30- or 90-min incubation. (E-F) A representative example of the CTV dilution (E) and calculated division index of OT-I T cells (F) after three days of stimulation, either without removing the stimulation (no wash) or by washing away IF/free pMHC after 30 or 90 min, respectively. Statistical significance was determined by two-way ANOVA on log-transformed data with post-hoc Sidak’s multiple comparison test. (B-C, F) *n =* 3 in three independent experiments. p-values are indicated in the Figure.

The importance of multivalency provided by the IF-pMHC^(SIIN)^ to enable stable interactions with the T cells became clear when OT-I T cells were washed after incubation with IF-pMHC^(SIIN)^ or free pMHC^(SIIN)^ (**Figure 2D**). Here, we found that only the IF-pMHC^(SIIN)^ were able to induce IFNγ production (**Figure S1F**) and substantial T cell proliferation (**Figure 2E-F**), demonstrating that robust multivalent interactions with the IFs are essential to initiate T cell activation. We hypothesize that these multivalent interactions will be even more relevant to enable prolonged T cell stimulation in vivo following intravenous (iv) injection, as this may prevent IFs from washing off from the T cells surface by shear forces in the bloodstream. Finally, as a result of the high affinity of the OT-I TCR for the SIINFEKL ligand, we observed that solely presenting pMHC^(SIIN)^ on the IFs appeared sufficient to stimulate OT-I T cells, and that co-presentation of αCD28, recombinant IL-2 or both did not further boost T cell activation (**Figure S1G**).

### 2.3 Activation and transcriptional characterization of mouse T cells stimulated ex vivo by IFs presenting lower affinity pMHC

We then continued to study the performance of IFs in a lower antigen affinity model, which more closely resembles anti-tumor T cell responses in cancer patients. To this end, we immobilized H-2Kb with the SIITFEKL peptide (pMHC^(SIIT)^), the T4 ligand for the OT-I TCR which has an approximate ten-fold lower affinity than the parental SIINFEKL ligand.^[24,25]^ When comparing IF-pMHC^(SIIT)^ with IFs that co-presented recombinant IL-2 on the same IFs (IF-pMHC^(SIIT)^/IL-2), we observed that both IF-pMHC^(SIIT)^ and IF-pMHC^(SIIT)^/IL-2 activated OT-I T cells, as indicated by co-expression of CD69 and CD25 (**Figure 3A**) and induction of IFNγ production (**Figure 3B**), while significantly outperforming their free counterparts in this lower affinity system. Interestingly, three days after stimulation, the co-presentation of IL-2 on the IFs proved to be highly favorable, as OT-I T cells stimulated with IF-pMHC^(SIIT)^/IL-2 were not only more viable (**Figure S2A**), but also proliferated significantly more compared to T cells stimulated with IF-pMHC^(SIIT)^ alone (an average of 2.35 cycles for IF-pMHC^(SIIT)^/IL-2 versus 0.51 cycles for IF-pMHC^(SIIT)^ in three days, **Figure 3C**). This beneficial effect on T cell proliferation was not observed for IF-pMHC^(SIIT)^ filaments co-presenting αCD28 (**Figure S2B**). Moreover, IF-pMHC^(SIIT)^/IL-2 also outperformed IF-pMHC^(SIIT)^ from a functional perspective, as IF-pMHC^(SIIT)^/IL-2 clearly induced OT-I T cells with higher target cell killing potential (**Figure 3D**). We conclude that co-presentation of IL-2 and pMHC^(SIIT)^ immobilized on IFs enhances the proliferation and functionality of T cells expressing a lower affinity TCR. Presenting IL-2 in close proximity to a TCR trigger on IFs could furthermore be beneficial to direct IL-2 specifically to T cells and prevent off-target binding and toxicity in vivo.^[17]^

**Figure 3.**
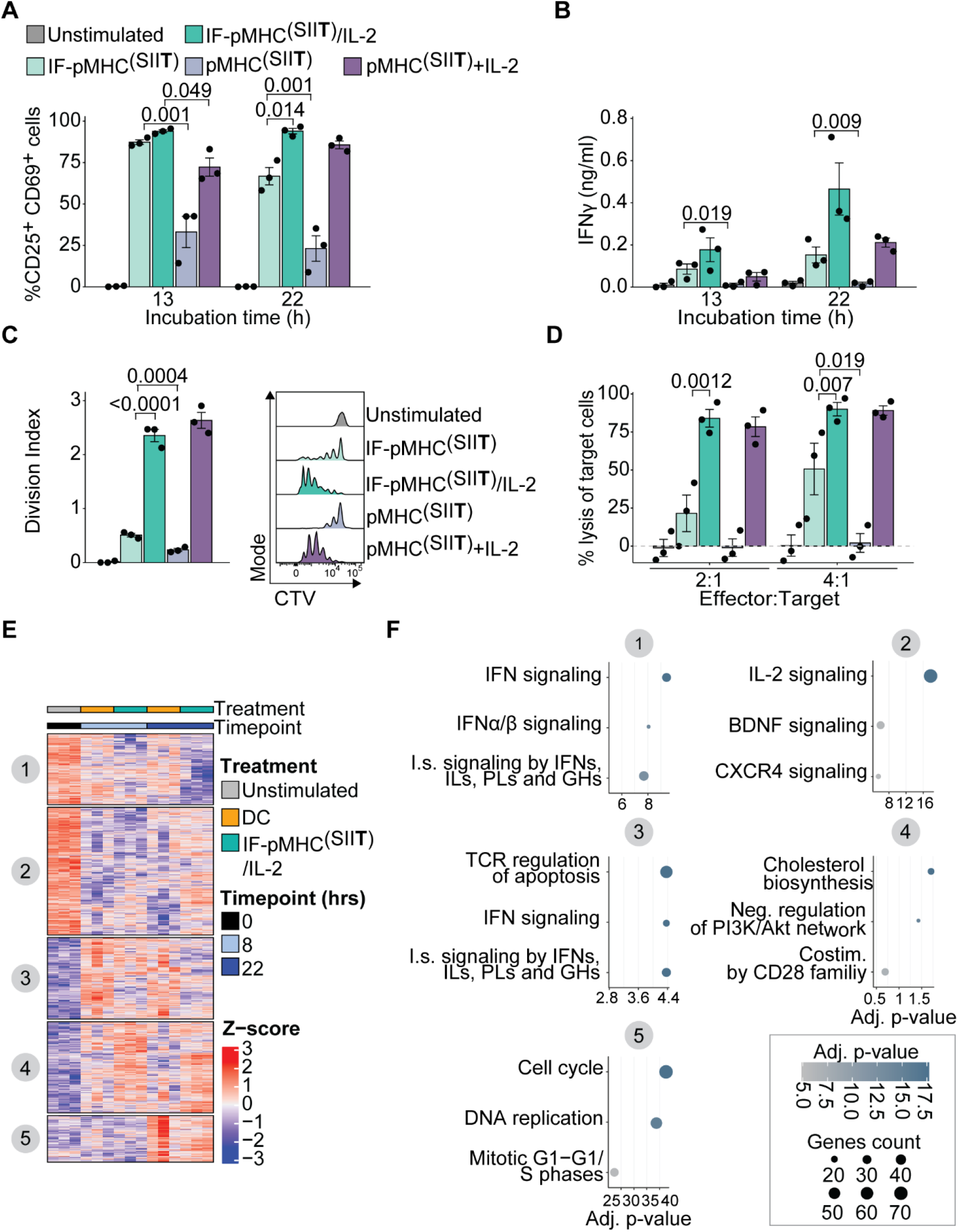
Activation and transcriptional characterization of mouse T cells stimulated by IFs presenting lower affinity pMHC. (A) Flow cytometry quantification of the percentage of activated CD25^+^CD69^+^ OT-I T cells at different time points. Statistical significance was determined with two-way ANOVA on logit-transformed data with post-hoc Tukey’s multiple comparison test. (B) The production of IFNγ by OT-I T cells at different time points by ELISA. Statistical significance was determined with two-way ANOVA on log-transformed data with post-hoc Tukey’s multiple comparison test. (C) OT-I T cells were stimulated for three days, after which the division index based on CTV dilution (right) was quantified as a measure of proliferation. Statistical significance was determined with one-way ANOVA on log-transformed data with post-hoc Tukey’s multiple comparison test. (D) Flow cytometry quantification of the percentage of lysed B16-OVA melanoma target cells 24 hours after co-incubation with OT-I T cells pre-stimulated for 20 hours. Statistical significance was determined with two-way ANOVA on logit-transformed data with post-hoc Tukey’s multiple comparison test. (A – D) *n =* 3 in three independent experiments. p-values are indicated in the Figure. (E-F) Antigen-specific OT-1 I T cells were either unstimulated (0 hours), or stimulated for 8 or 22 hours with SIITFEKL-loaded DCs or IF-pMHC^(SIIT)^/IL-2 in triplicate in one experiment. (E) Heatmap depicting z-scores of differential genes (Benjamini-Hochberg adjusted p-value <0.05 and fold change > 2-fold) compared to unstimulated cells (0 hours). Gene clusters were obtained using k-means clustering. The columns are ordered according to treatment timepoint and mouse (three biological replicates per condition). (F) Gene ontology analysis of the five gene clusters as depicted in (E), for which the three most enriched terms are visualized. X-axis and colors depict -log10 of the Benjamini-Hochberg-adjusted p-values for overrepresentation of the most enriched gene ontology terms in each cluster compared to expected. Higher values mean stronger enrichment for the specific terms. Dot size indicates how many genes are present in the enriched gene sets. IFN: Interferon; I.s.: Immune system; IL: Interleukins; PL: Prolactin; GH: growth hormones; BDNF: brain-derived neurotrophic factor; CXCR4: C-X-C Motif Chemokine Receptor 4; Neg. regulation: Negative regulation; Costim.: Costimulation; Adj.: Adjusted.

Next, we studied to what extent IFs resemble natural DCs in their stimulation of antigen-specific T cells. To this end, we molecularly characterized the transcriptional programs underlying T cell activation following antigen-specific OT-I T cell stimulation with IF-pMHC^(SIIT)^/IL-2, or with primary murine DCs pulsed with OVA protein. Both IFs and DCs induced similar expression of early activation markers CD69 and CD25 in OT-I T cells (**Figure S3A-B**). We then performed bulk RNA-sequencing of unstimulated OT-I T cells, and of OT-I T cells after 8 and 22 hours of stimulation. After filtering of lowly expressed genes, we reproducibly quantified approximately 11,000 genes in all samples. To assess how exposure of T cells to either IFs or DCs impacted their respective transcriptomes, we identified differentially expressed genes relative to the untreated control at 0 hours. Hierarchical clustering of these ∼1600 genes revealed five major clusters (**Figure 3E**). For each cluster, we performed gene ontology analyses to classify the biological processes (**Figure 3F**). Both DCs and Ifs stimulation activated cell cycle and cell division gene expression programs after 22 hours, highlighting that similar pathways were induced and validating the activating potential of the IFs (Figure 3E-F, cluster e). Although in general very similar pathways were induced by DCs and IF, on a more detailed level we observed that T cells stimulated with IF-pMHC^(SIIT)^/IL-2 specifically upregulated genes related to IL-2 signaling at 22 hours (Figure 3E-F, clusters b and d). We furthermore found that IFs stimulation evoked lower interferon signaling compared to T cells that were activated by DCs (Figure 3E-F, clusters a and c). One-on-one comparison of gene expression at 8 and 22 hours after stimulation versus unstimulated OT-I T cells demonstrated that DC and IF-pMHC^(SIIT)^/IL-2 stimulation induced a similar number of upregulated and downregulated genes (**Figure S3C-D**). Together, these data indicate that both IF-pMHC^(SIIT)^/IL-2 and DC stimulation of T cells activate similar transcriptional profiles associated with proliferation, whereas T cells respond differently with respect to their IL-2 and IFN signaling.

### 2.4 Immunofilaments effectively stimulate NY-ESO-1-specific CD8+ T cells and human CD19 CAR T cells ex vivo

Next, we evaluated the versatility of the IF platform by testing their ability to activate and expand other types of antigen-specific T cells beyond the murine OT-I T cell system, including a humanized transgenic T cell model and a human CAR T cell system. We conjugated recombinant IL-2 onto IFs together with a human HLA-A2.1-NY-ESO-1157-165 (SLLMWITQV). We used these IFs (IF-A2^(NY-ESO-V)^ and IF-A2^(NY-ESO-V)^/IL-2) to stimulate transgenic murine CD8^+^ T cells that express the cognate 1G4 TCR specific for the human NY-ESO-1157-165 (**Figure 4A**).^[26]^ The 1G4 T cells could be activated by IFs presenting A2^(NY-ESO-V)^ alone, but they displayed a remarkably enhanced production of IFNγ (**Figure 4B**) and proliferation (**Figure 4C, D**) when IL-2 was co-presented (IF-A2^(NY-ESO-V)^/IL-2). These data confirm our previous observations indicating that immobilized IL-2 can enhance CD8^+^ T cell responses when it is co-presented with pMHC on IF.

**Figure 4.**
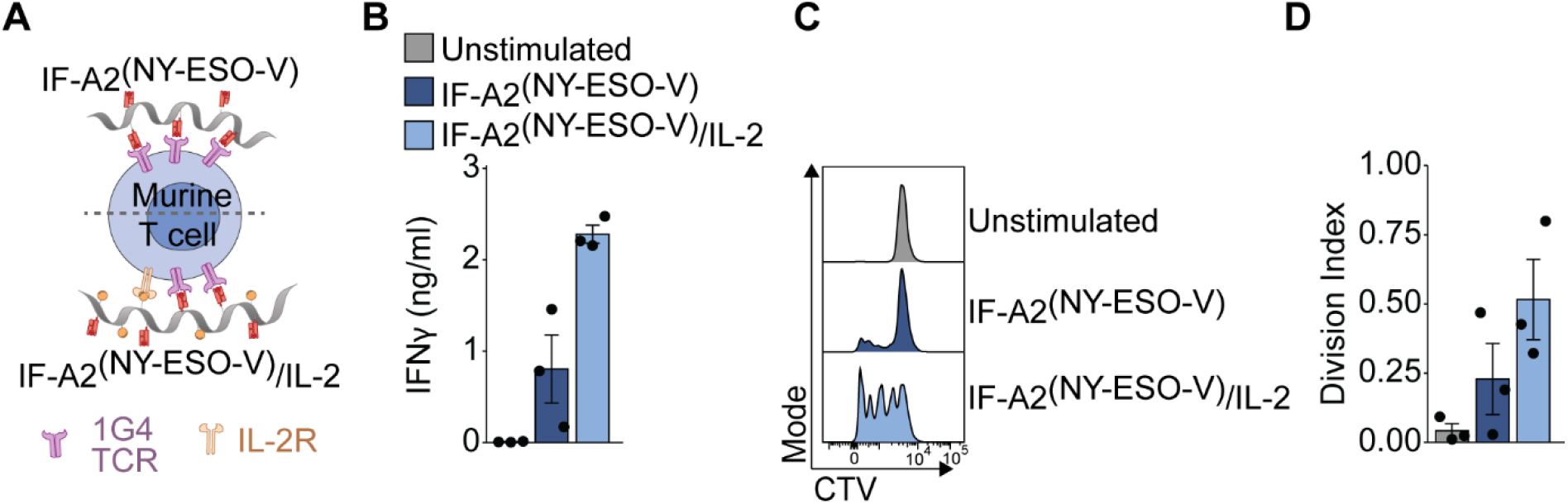
Immunofilaments effectively stimulate NY-ESO-1-specific murine CD8+ T cells. (A) Schematic overview of IF-A2^(NY-ESO-V)^ and IF-A2-^(NY-ESO-V)^/IL-2 for stimulation of murine 1G4 T cells. (B) IFNγ production of 1G4 T cells after 24 hours of stimulation with IF-A2^(NY-ESO-V)^ and IF-A2-^(NY-ESO-V)^/IL-2. Statistical significance was tested with an unpaired t-test on log-transformed data. (C, D) A representative example of the CTV dilution (C) and calculated division index of 1G4 T cells after three days of stimulation. Statistical significance was tested with an unpaired t-test on log-transformed data. *n =* 3 in three independent experiments.

Next, we stimulated human CD8^+^ T cells transfected with a transgenic TCR against NY-ESO-1 (**Figure S4A**), and we observed an increase in their expression of CD25 (**Figure S4B**). We furthermore found that only TCR-transfected but not mock-transfected human T cells produced IFNγ and proliferated in response to IFs (**Figure S4D-F**), demonstrating the antigen-specificity of the IF.

Artificial APCs are used frequently for the bulk ex vivo expansion of T cells genetically engineered to express CARs, after which these potentiated T cells are infused back into patients for immunotherapeutic purposes. CAR T cell expansion usually relies on αCD3/αCD28-coated Dynabeads for T cell stimulation. Although this polyclonal stimulation can efficiently expand CAR T cells, bystander non-CAR T cells are expanded at equal rates which could pose threats with respect to unwanted side effects.^[27]^ We therefore probed the potential of the IFs to selectively expand CD19-directed CAR T cells by activating them through engagement of the CARs. We produced IFs that present Protein L (ProL), which binds to the variable kappa light chains of immunoglobulins and thereby specifically crosslinks any CAR on the T cell surface to induce T cell activation.^[28,29]^ In addition, we prepared IFs that co-presented ProL and anti-human CD28 or recombinant IL-2 (**Figure 5A**). A mixture of CD4^+^ and CD8^+^ T cells were lentivirally-transduced with a second generation CD19 CAR construct.^[30]^ After expansion and resting, the CD19 CAR T cells were re-stimulated with IF. We observed increased IFNγ production by CAR T cells stimulated with IF-ProL/IL-2 compared to those stimulated with IF-ProL or IF-ProL/αCD28 (**Figure 5B-D**), resulting in a 9.3- and 5.0-fold enhanced IFNγ levels on day 5, respectively. Whereas αCD28 did not seem to have any beneficial effect on CAR T cell activation, the co-presentation of ProL with IL-2 synergized to boost CAR T cell stimulation (**Figure S5**), again demonstrating that co-immobilizing and TCR-triggering molecule and IL-2 enhances T cell prolfieration.

**Figure 5.**
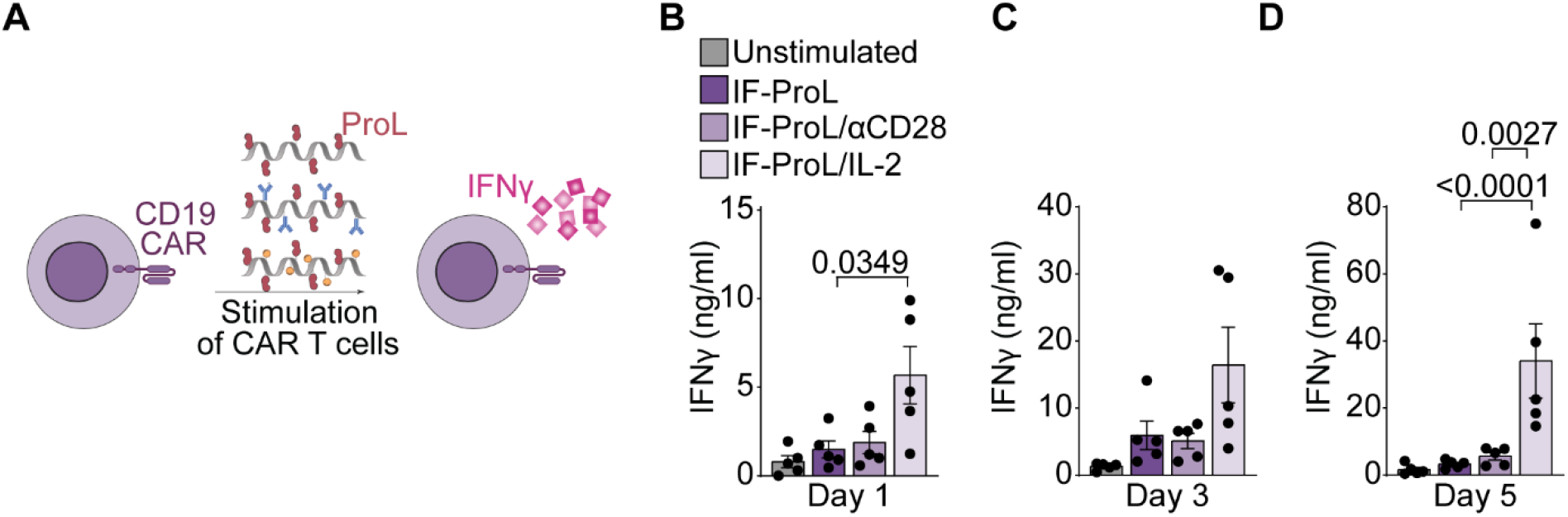
Immunofilaments effectively stimulate human CD19 CAR T cells ex vivo. (A) Schematic overview of IF-ProL, IF-ProL/αCD28 and IF-ProL/IL-2 for the stimulation of human T cells lentivirally-transduced with a CD19 CAR construct. (B-D) IFNγ production after stimulation with IFs for 1 day (B), 3 days (C) and 5 days (D). Statistical significance was determined by two-way ANOVA on log-transformed data with post-hoc Tukey’s multiple comparison test. *n =* 5 in five independent experiments. p-values are indicated in the Figure.

Together, these data exemplify the versatility of IFs in their ability to effectively activate different types of antigen-specific T cells with different affinities and genetically engineered T cells. As such, IFs constitute a modular platform for ex vivo expansion of T cells for which the presented biomolecules can be easily adjusted to the requirements of the system.

### 2.5 In vivo biodistribution of IFs following intravenous administration

Before applying IFs therapeutically for in vivo expansion of T cells, we assessed their biodistribution after iv administration to analyze whether they could reach lymphoid organs. We labeled non-functionalized IFs and IF-pMHC^(SIIN)^ with ^111^In and injected them iv into mice. Both non-functionalized IFs and IF-pMHC^(SIIN)^ were detectable in the blood for up to 24 hours after administration though IF-pMHC^(SIIN)^ was cleared from the systemic circulation slightly faster (6.3% ID/g for non-functionalized IFs versus 3.8% ID/g for IF-pMHC^(SIIN)^, **Figure 6A**). Both IFs displayed similar biodistribution patterns across the organs after 24 hours, with substantial accumulation in spleen, liver, lungs and kidneys (**Figure 6B-C**). In addition to the significant accumulation of non-functionalized IFs and IF-pMHC^(SIIN)^ in the spleen, we also observed notable amounts of IFs in other secondary lymphoid organs in which T cells predominantly reside, including the popliteal (Figure 6B), the axillary and inguinal lymph nodes (**Figure 6D**). This facilitates the interaction of IFs with antigen-specific T cells, which is essential to trigger T cell responses in vivo.

**Figure 6.**
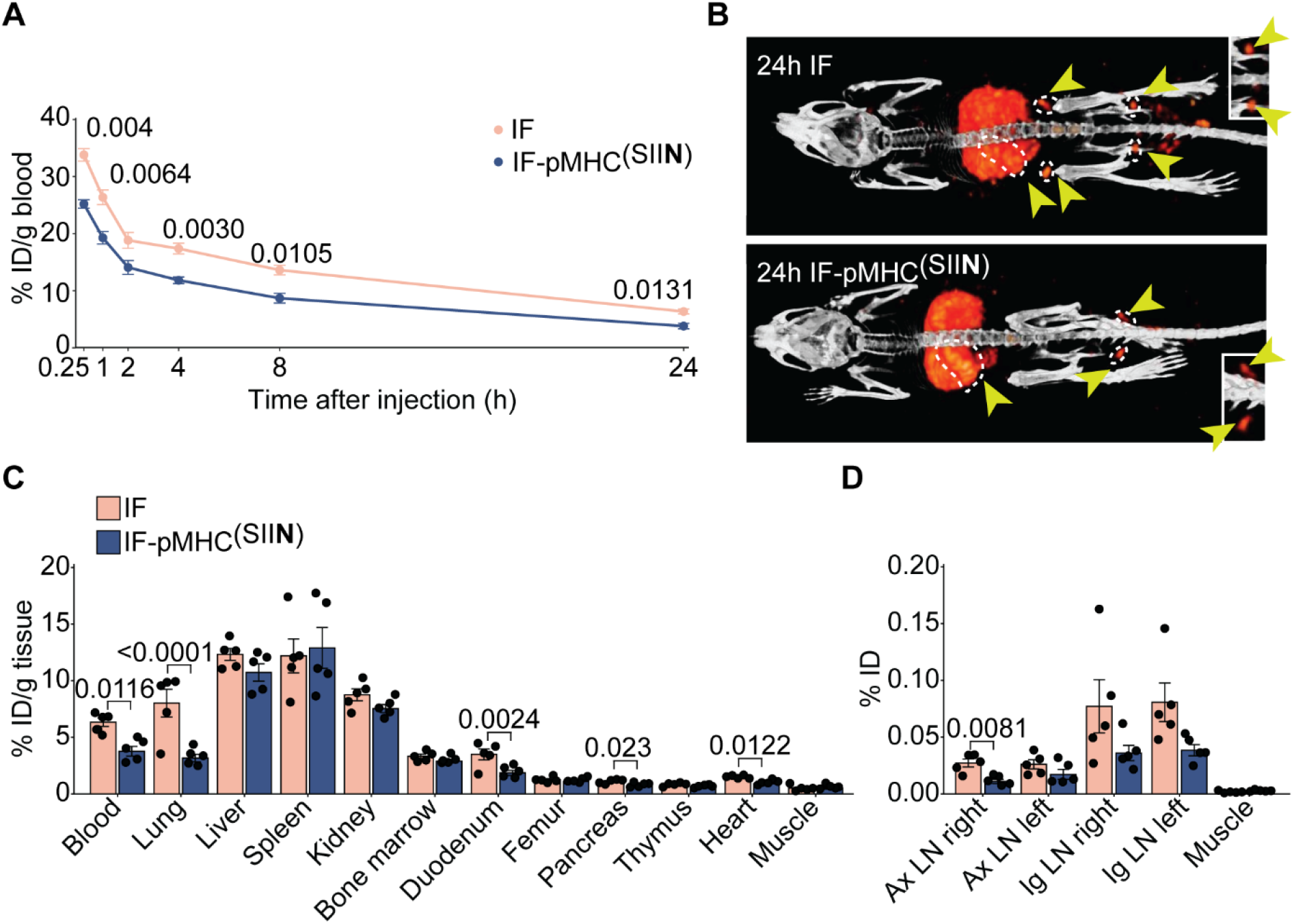
The in vivo biodistribution of IFs after intravenous injection. (A) Kinetics of ^111^In-labelled IFs and IF-pMHC^(SIIN)^ in the blood after iv injection in WT C57BL/6 mice, expressed as % of injected dose (ID) per gram of blood. *n = 6* for t = 0.25, 1, 2, 4 hours and *n = 5* for t = 8 and 24 hours in one independent experiment. Statistical significance was determined with mixed effect analysis and post-hoc Sidak’s multiple comparison test, p-values are indicated in the Figure. (B) 2D maximum intensity projection SPECT/CT images 24 hours after iv injection of ^111^In-labeled IFs and IF-pMHC^(SIIN)^. Yellow arrows indicate secondary lymphoid organs (dotted white line indicates spleen and lymph nodes). Inset: Magnification of the popliteal lymph nodes. (C) Quantitative ex vivo analysis of the biodistribution of IFs 24 hours after iv injection across different organs expressed as % of ID per gram of tissue. Statistical significance was determined by two-way ANOVA on log-transformed data with post-hoc Sidak’s multiple comparison test. *n = 5* in one independent experiment. p-values are indicated in the Figure. (D) Quantitative ex vivo analysis of the % of the ID of IFs 24 hours after iv injection in the axillary lymph nodes (Ax LN) and inguinal lymph nodes (Ig LN), compared to a piece of quadriceps muscle with the same weight. Statistical significance was determined by two-way ANOVA on log-transformed data with post-hoc Sidak’s multiple comparison test. *n = 5* in one experiment. p-values are indicated in the Figure.

To investigate whether the biodistribution of IF-pMHC^(SIIN)^ would be affected by the presence of antigen-specific T cells in vivo, we administered ^111^In IF-pMHC^(SIIN)^ iv to mice that received adoptively transferred WT T cells or OT-I T cells. The biodistribution of IF-pMHC^(SIIN)^ was not altered by the presence of antigen-specific OT-I T cells, apart from a slightly higher accumulation of IF-pMHC^(SIIN)^ in the spleen, which may suggest that IF-pMHC^(SIIN)^ interact with and are retained in the spleen by OT-I T cells (**Figure S6**). Taken together, these data demonstrate that non-functionalized IFs and IF-pMHC^(SIIN)^ have similar biodistribution patterns, are available for >24 hours in the blood and can reach lymphoid organs.

### 2.6 Immunofilaments induce antigen-specific T cell proliferation in vivo

The observation that IFs readily reach lymphoid organs upon administration in vivo (Figure 6) provided us with a unique opportunity to exploit IFs for direct T cell expansion in vivo. To this end, we adoptively transferred unstimulated OT-I T cells into mice and 24 hours later injected IF-pMHC^(SIIN)^ or non-functionalized IFs mixed with free pMHC^(SIIN)^ (**Figure 7A**). The IF-pMHC^(SIIN)^ induced proliferation of OT-I T cells in vivo to a much higher extent than when these two components were given separately **(Figure 7B-C**). These findings confirm the importance of providing T cell-activating biomolecules on a polymer backbone to facilitate docking to T cells and support downstream signaling by multivalency, probably combined with the effect of an more favorable half-life and biodistribution for the IF-pMHC^(SIIN)^. When IF-pMHC^(SIIN)^ were administrated to mice that received adoptive transfer of wildtype CD8^+^ T cells instead of antigen-specific OT-I T cells, we did not observe any proliferation (**Figure S7A-C**). These findings clearly demonstrate that IF-pMHC^(SIIN)^ trigger T cell activation in an antigen-specific manner in vivo. Furthermore, IF-A2^(NY-ESO-V)^/IL-2 were also able to trigger proliferation of antigen-specific 1G4 T cells what were adoptively transferred into recipient mice (**Figure S8**), showing that this observation also holds true for other antigens and other T cells expressing different TCRs.

**Figure 7.**
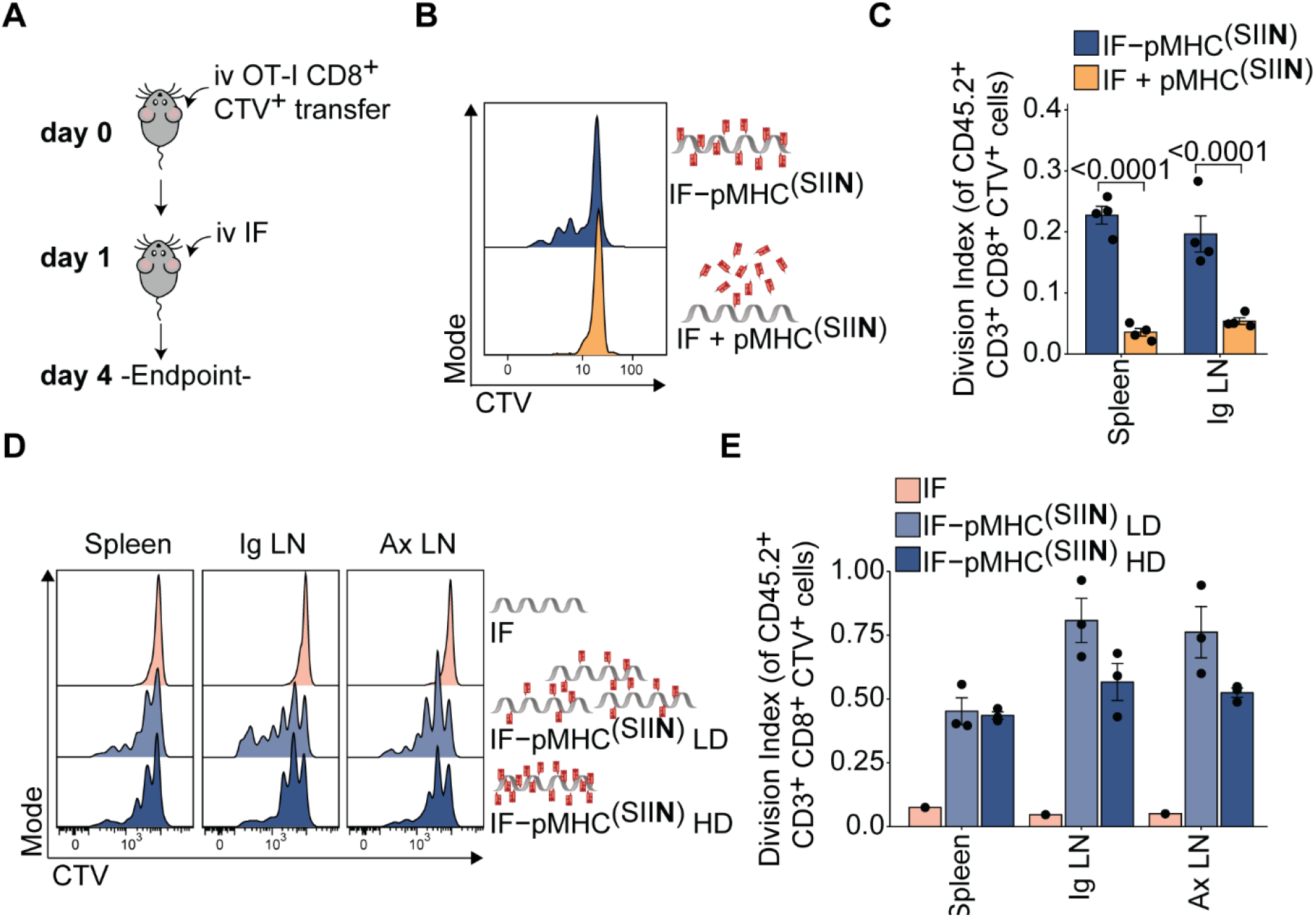
Immunofilaments expand antigen-specific T cells in vivo. (A) Schematic overview of experimental setup. (B-C) Ex vivo flow cytometric analysis of CTV dilution (B) and quantification (C) of the division index of CD45.2^+^CD3^+^CD8^+^ OT-I T cell in the spleen 3 days after iv administration of 0.29 μg free pMHC^(SIIN)^ + IFs or IF-pMHC^(SIIN)^ in CD45.1^+^ WT C57BL/6 adoptively transferred with 5×10^5^ OT-I T cells. Statistical significance was determined on log-transformed data with two-way ANVOA and post-hoc Dunnet’s multiple comparison test. *n = 3* in one independent experiment. (D-E) Ex vivo flow cytometric analysis of CTV dilution (D) and quantification (E) of the division index of OT-I T cell in the spleen and inguinal (Ig) and axillary (Ax) lymph nodes 4 days after iv administration of IFs alone or IF-pMHC(SIIN) with a low density (LD): ∼5 pMHC per IF) or high density (HD): ∼16 pMHC per IF). The total amount of pMHC^(SIIN)^ (1 μg) that was administered was kept constant. Statistical significance was tested with two-way ANOVA on log-transformed data with Sidak’s multiple comparison test. *n = 3* in one independent experiment.

To investigate how IFs design affects T cell proliferation, we next compared proliferation of adoptively transferred OT-I T cells after injection of IF-pMHC^(SIIN)^ decorated with different numbers of pMHC^(SIIN)^. Keeping the amount of injected pMHC^(SIIN)^ constant, we varied the amount and spacing of pMHC^(SIIN)^ per polymer (low density (LD): ∼5 pMHC per IF, high density (HD): ∼16 pMHC per IF) and thus varied the total amount of IFs that we administered to the animals. As we observed that the non-functionalized IFs backbone itself does not induce OT-I T cell proliferation (**Figure 7D-E**), this allowed us to study the impact of the pMHC^(SIIN)^ density on IFs in vivo. We observed a slightly higher proliferation of OT-I T cells upon administration of LD IF-pMHC^(SIIN)^ compared with HD IF-pMHC^(SIIN)^ in the lymph nodes but not in the spleen, which indicated that a lower density of pMHC^(SIIN)^ is sufficient for T cell activation (Figure 7D-E). We hypothesize that a low density of pMHC^(SIIN)^ already being effective is the direct consequence of the administration of higher numbers of IFs molecules in total, thereby increasing the chances of IF-pMHC^(SIIN)^ to meet and interact with antigen-specific T cells in vivo. Next, we studied the importance of the time interval between adoptive cell transfer of OT-I T cells and injection of IF. We found that for both LD and HD IFs the resulting T cell proliferation was similar irrespective of whether IF-pMHC^(SIIN)^ were administrated 10 min, 4 hours or 24 hours after adoptive cell transfer (**Figure S7D-G**). Finally, we studied how the number of adoptively transferred OT-I T cells and the dose of IF-pMHC^(SIIN)^ affects T cell proliferation in vivo after iv administration of IFs (**Figure S7H**). Whereas the number of transferred OT-I T cells did not affect their proliferation, we observed a clear effect of the IF-pMHC^(SIIN)^ dose by iv injection on the level of OT-1 proliferation both in terms of the percentage of proliferating cells and their division index (**Figure S7I-J**). Within the tested dose range, we did not observe dose-dependent proliferation when IFs were administered subcutaneously (sc) as there was high proliferation for all doses tested, suggesting that sc delivery of IFs is highly efficient (**Figure S7K-M**). In conclusion, IFs are able to stimulate antigen-specific T cells in vivo through iv and sc administration, which leads to T cell expansion.

### 2.7 Immunofilaments inhibit primary tumor growth and reduce metastatic spread in vivo

Finally, we investigated the potential of IFs in vivo in the context of cancer models. We made use of a well-established aggressive melanoma model in which the effect of IFs on the establishment of pseudometastasis can be probed. OT-I T cells were adoptively transferred into mice and stimulated through iv administration of IF-pMHC^(SIIN)^ or free pMHC^(SIIN)^ (**Figure 8A**). On day five, B16-OVA melanoma cells were injected iv into the tail vein to induce the development of metastases in the lungs and 19 days post-injection, lungs were isolated and pulmonary lesions enumerated (**Figure 8B**). Although the transfer of OT-I T cells alone modestly reduced the number of metastatic lesions in the lungs (an average of 162 metastatic nodules), iv treatment with IF-pMHC^(SIIN)^ significantly decreased the number of metastatic nodules to an average of 97 (**Figure 8C**) compared to PBS treatment alone. The administration of free pMHC^(SIIN)^ did not lead to a reduction in the number of metastatic lesions compared to the controls, which is probably the result of fast clearance and poor OT-I T cell stimulation. We next examined the lungs to establish the total metastatic burden, which we defined as the percentage of the lung area occupied by metastatic lesions. This analysis does not only consider the number of metastatic lesions but also incorporates the size of the individual nodules, and we again found that IF-pMHC^(SIIN)^ significantly outperforms pMHC^(SIIN)^ in reducing the metastatic burden (**Figure 8D**).

**Figure 8.**
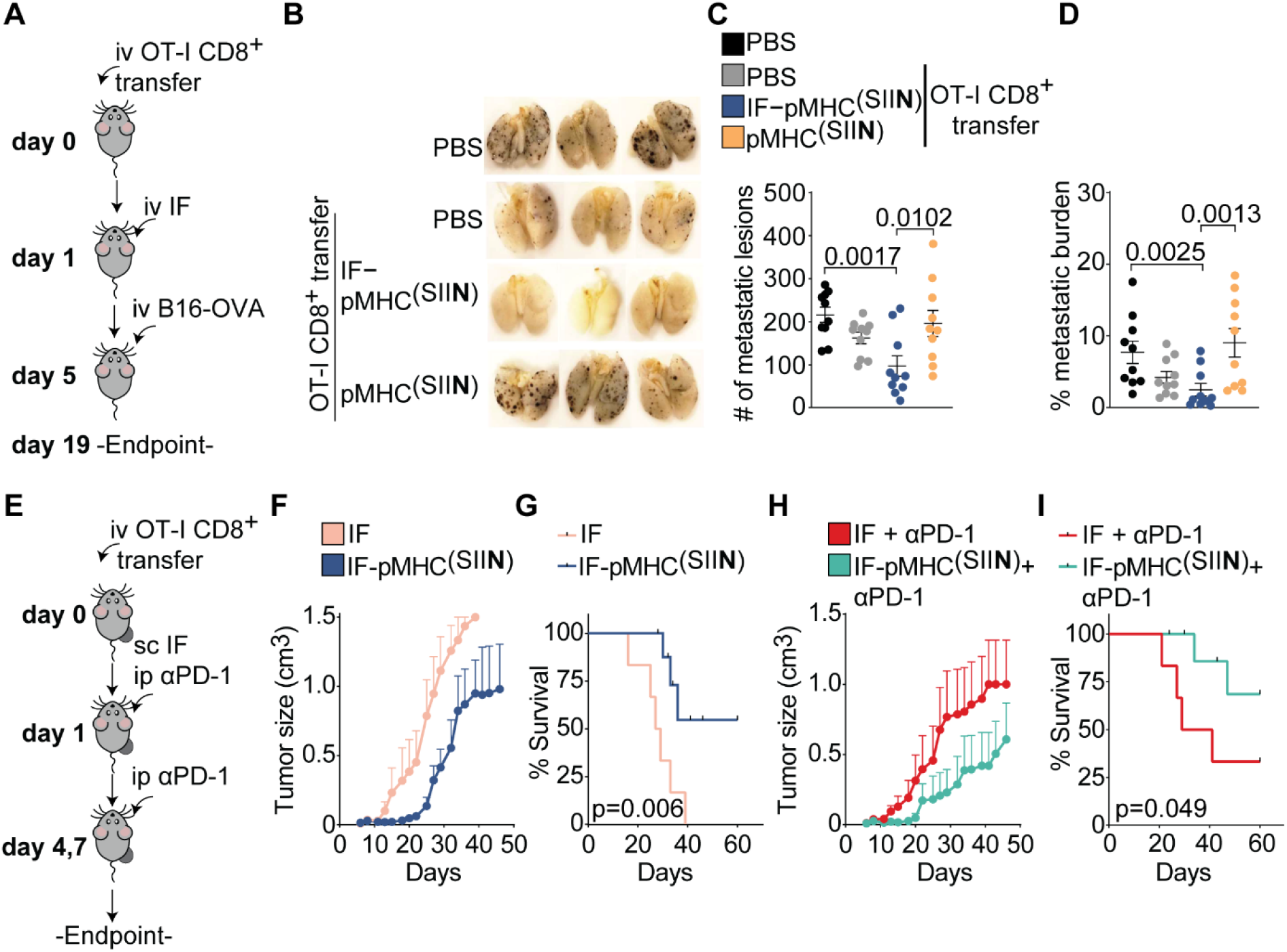
In vivo administration of IFs inhibits primary tumor growth and prevents development of metastases. (A) Overview of experimental setup of the lung metastasis model. OT-I T cells were adoptively transferred into WT mice, followed by a single iv injection of IFs on day 1. On day 5, B16-OVA tumor cells were injected iv and on day 19 the lungs were collected and fixed overnight in Fekete’s solution. The number of metastatic nodules in the lungs was enumerated. (B-C) Representative overview of metastatic lesions (in brown/black) in the lungs at day 19 (B) and enumeration of the total number of metastatic nodules in the lungs (C). *n =* 10 in 2 independent experiments. Statistical significance was determined with one-way ANOVA and post-hoc Dunnet’s multiple comparison test. (D) Quantification of the metastatic burden of the lungs on day 19, defined as the percentage of the lungs occupied by metastatic lesions. Statistical significance was determined by one-way ANOVA on logit-transformed data and post-hoc Dunnet’s multiple comparison test. (E) Overview of the experimental setup of the sc injection of IF. OT-I T cells were adoptively transferred into B16-OVA tumor-bearing mice, followed by a single sc injection of IFs on day 1 with or without three times intraperitoneal injection of αPD-1. (F-G) Quantification of the size of B16-OVA tumors (F) and the survival (G) of mice treated with non-functionalized IFs or IF-pMHC^(SIIN)^. *n =* 6-9 in one independent experiment. (H-I) Quantification of the size of B16-OVA tumors (H) and the survival (I) of mice treated with IFs + αPD-1 or IF-pMHC^(SIIN)^ + αPD-1. (G, I) Statistical significance was determined with the Gehan-Breslow-Wilcoxon test. Mice that did not reach the tumor size of 1500 mm^3^ were marked as censored. *n =* 6-9 in one experiment.

To investigate how IFs perform in a therapeutic cancer model where an established tumor is present, we adoptively transferred OT-I T cells into mice bearing B16-OVA melanoma tumors in their flank. Next, a single dose of IF-pMHC^(SIIN)^ was administrated sc in close proximity to the established tumors (**Figure 8E**). Mice receiving a single dose of IF-pMHC^(SIIN)^ displayed a substantial delay in tumor growth and a significantly enhanced survival compared with mice that received IFs alone (**Figure 8F-G, Figure S9A-E**). We next studied if stimulation and proliferation of antigen-specific T cells with IF-pMHC^(SIIN)^ could synergize with alleviating immune suppression through blocking of the immune checkpoint PD-1 on T cells. To this end, we combined sc administration of IF-pMHC^(SIIN)^ with intraperitoneal injections of αPD-1. The results confirmed that this combination provides a clear benefit in terms of delaying tumor growth and significantly improving survival compared to treatment with αPD-1 alone (**Figure 8H-I**). Histological analysis of the T cell infiltration in these tumors showed that mice that were treated with either IF-pMHC^(SIIN)^ or IF-pMHC^(SIIN)^ + αPD-1 had slightly elevated numbers of CD8^+^ T cells infiltrating in their tumors (**Figure S9F-J**). These results clearly indicate that OT-I T cells stimulated by a single dose of IFs are highly functional and migrate to the tumor where they can combat the growth of primary B16-OVA tumors in vivo.

Altogether, our findings demonstrate that nanosized IFs constitute a powerful new tool that elicits strong T cell activation ex vivo and in vivo, resulting in potent anti-tumor efficacy. Previously, other aAPCs have been developed that can be administered in vivo to stimulate T cells, but these aAPC platforms predominantly consist of microsized beads.^[31–35]^ Their large surface area and low surface curvature promotes interaction with the T cell surface, resulting in strong T cells responses that cannot be obtained with nanosized beads.^[36,37]^ However, their micrometer range size prohibits effective intravenous injection as they are rapidly cleared by the reticuloendothelial system, have limited access to lymphoid organs and can pose significant safety concerns.^[36,37]^ Instead, IFs provide a unique alternative as they are nanosized and at the same time give rise to strong T cell activation and initiate proliferation programs. We hypothesize that this is due to the semiflexible nature of the filaments, which supports receptor clustering on the cell surface to drive signaling when biomolecules are presented multivalently to T cells. Consequently, our nanosized IFs do not suffer from the challenges associated with iv delivery of microsized beads, but instead remain circulating in the blood for at least 24 hours following iv administration and can effectively reach lymphoid organs. These IFs effectively expand antigen-specific T cells in vivo that acquire tumor-killing capacities and are able to migrate, thereby delaying tumor growth and extending survival. We furthermore demonstrate IFs can be applied both for iv and sc injection, putting forward IFs as a novel, modular and broadly applicable platform adding to the arsenal of immunotherapies to fight cancer.

## 3. Conclusion

We have developed a nanosized IF platform able to present pMHC, costimulatory molecules and cytokines in a multivalent manner, which effectively stimulate antigen-specific T cells. Our findings demonstrate that nanosized IFs elicit strong T cell activation ex vivo and initiate transcriptional programs largely similar to those evoked by natural DCs. We furthermore observed that co-immobilization of IL-2 and pMHC on IFs evokes particularly potent T cell responses for lower affinity T cell antigens. These IFs constitute a versatile platform that can be applied for the ex vivo expansion of antigen-specific T cells or CAR T cells. The most important benefit of these nanosized IFs is that owing to their small size, they can readily be applied to activate and expand T cells in vivo through iv and sc injection. After iv injection, IFs are able to reach lymphoid organs and evoke strong T cell responses in vivo. As a result, even a single dose of IFs has a remarkable antitumor effect, as it inhibits metastases formation in a melanoma model and reduces primary tumor growth in synergy with immune checkpoint blockade. We conclude that these nanosized IFs are a novel, powerful and versatile type of aAPC. They constitute a major step forward in the treatment of cancer as in vivo administration of IFs may not only specifically activate and expand existing tumor specific T cells or TIL, but may also T cells or CAR-T cells revitalize previously administered by ACT, thus enhancing their lifespan and therapeutic efficacy.

## 4. Experimental Section

### Immunofilament synthesis and characterization

IFs were prepared as described before.^[17,19,20]^ Briefly: isocyanopeptide monomers with non-functional methoxy and functional azide groups were polymerized in a 30:1 ratio using a nickel catalyst (1/10,000 ratio), yielding polymers with statistically one azide group every 3.5 nm.^[16,18,21]^ Next, 60% of the azides were reacted with DBCO-PEG4-biotin according to literature procedures.^[22]^ The average length of the azide/biotin polymers was determined using atomic force microscopy (AFM, nanoscope III, digital instruments), operated in tapping mode in air. The polymers were dissolved in MilliQ (10 μg/mL) and drop casted on freshly cleaved mica for 5 minutes, after which the sample was dried under a nitrogen flow. From the resulting images the polymer length was determined using ImageJ.^[38]^ The average length determined was 407 nm ± 207 nm, calculated from 161 values.

### MHC production

MHC were prepared as described.^[39]^ The constructs for the heavy chains (HLA-A2, H-2Kb) and human beta-2-microglobulin (hβ2m) were generously provided by M. Toebes and T.N. Schumacher from the NKI. They were produced as inclusion bodies in E.coli BL21(DE3)pLysS using T7 RNA polymerase/promoter system.^[40]^ Isolated inclusion bodies were solubilized in denaturing buffer (8 M urea/100 mM Tris•Cl, pH 8). Hβ2m was pre-folded in dialysis against 10 mM Tris•Cl (pH 7) in PBS. To prepare the final MHC complex, hβ2m and heavy chain were dissolved to final concentrations of 6 mM and 3 mM respectively in folding buffer (100 mM Tris•Cl, pH 8; 400 mM L-arginine; 2 mM EDTA; 5% glycerol; 5 mM reduced glutathione; 0.5 mM oxidized glutathione; Protease Inhibitor Cocktail, Roche Diagnostics) with 60 mM template peptide (NY-ESO-V: SLLMWITQV, which is a higher affinity analogue of the natural NY-ESO-1157-165 peptide where the terminal cysteine (C) amino acid is replaced by valine (V),^[41]^ SIINFEKL, SIITFEKL; GenScript). The folding reaction was incubated in 10 °C for 5 days. After filtration, concentration and buffer change to PBS, the complexes were purified via Size Exclusion Chromatography using HiLoad 16/600 Superdex 75pg column (Cytiva). Ready MHC was analyzed using SDS-PAGE and NanoDrop, concentrated, snap-frozen and stored at -80°C until further use.

### Protein functionalization

Proteins were functionalized using previous reported protocols.^[9,17]^ Briefly, pMHC^(SIIN)^, pMHC^(SIIT)^ and A2^(NY-ESO-V)^ were obtained from the refolding protocol, typically 0.5-2 mg/mL in PBS (pH 7.4). Mouse αCD28 was obtained from BioXcell, IL-2 was obtained from ProSpec and ProL was obtained from Acrobiosystems. Before use, mouse αCD28 and ProL were washed with borate buffer (pH 8.4, 50 mM) using 30 kDa spinfilters from Amicon, typically to 2-3 mg/mL. IL-2 was first reconstituted in MilliQ (10 mg/ml) and after 10-30 min further diluted with PBS (pH 7.4), typically to 1 mg/mL. To functionalize the proteins, DBCO-PEG4-NHS (Click chemistry tools, 100 mM in DMSO) and dye-NHS (10 mM in DMSO) were added to the protein stock solutions, typically 3-4 eq DBCO and 2.5-3 eq dye were used. Dyes used in this study are: AlexaFluor350-NHS (Thermo Fisher Scientific), Atto488-NHS (AttoTec), AlexaFluor594-NHS (Thermo Fisher Scientific) and AZdye647-NHS (click chemistry tools). For proteins in borate buffer, reactions were run for 2-3 hours at 4 °C. Proteins in PBS were left to react for 4-6 hours at 4 °C. After the reaction the proteins were purified using spin filtration against PBS with spinfilters of appropriate size (Amicon). The protein conjugates were analyzed with NanoDrop (Thermo Fisher Scientific), using the following extinction coefficients: MHCs (95000 M^-1^ cm^-1^), mouse αCD28 (210000 M^-1^ cm^-1^), ProL (35760 M^-1^ cm^-1^), IL-2 (11900 M^-1^ cm^-1^), DBCO (12000 M^-1^ cm^-1^), AlexaFluor350 (19000 M^-1^ cm^-1^), Atto488 (90000 M^-1^ cm^-1^), AlexaFluor594 (120000 M^-1^ cm^-1^) and AZdye647 (270000 M^-1^ cm^-1^). Data was measured at 280 nm (protein), 309 nm (DBCO) and the wavelength of the dye. The raw data was corrected for spectral overlap between the different components using correction factors as described.^[17]^ Typical obtained degrees of labelling (DOL) for DBCO was 0.5-3 and 0.5-2 for dyes.

### Immunofilament-protein conjugates

The DBCO/dye functionalized proteins were coupled to the IFs using a protocol as described before.^[9,17]^ Briefly, 100-200 μL of a stock solution of 1 mg/ml biotin/azide polymer was reacted with the required amount of protein in PBS (pH 7.4), typically 0.2-1 eq of protein irt free azides was used. All reactions were carried out in non-stick Eppendorf tubes (Fisher Scientific), with a final concentration of 0.2-0.25 mg/mL polymer. Reactions were first mixed for 4-5 hours at RT, followed by incubation at 4 °C overnight. Purification was performed following a literature protocol, using monoavidin resin (Thermo Fisher Scientific).^[22]^ Per 100 μg of polymer, 1 mL of monoavidin resin was used. After 2x washing of the resin with PBS (pH 7.4), the resin was added to the reaction mixtures and incubate for 1.5-2 hours at 4 °C. Next, the resin was washed with 1x PBS-tween (0.1%) and 4-5x PBS. After washing, a solution of 2mM biotin in PBS was added (300-400 μL) to elute the polymer bioconjugates from the monoavidin resin (1-2 hours incubation at 4 °C). Concentrations of the conjugated proteins were determined using fluorescence (Tecan spark 10M plate reader), using the soluble labelled proteins as the standard. Polymer concentration was determined using circular dichroism spectroscopy (JASCO J-810). With these concentrations the spacing/density of the protein on the polymer could be calculated (**Table 1, Table 2**).

### Preparation of ^111^In-immunofilaments

To label IFs with ^111^In, first diethylenetriaminepentaacetic acid (DTPA)-PEG4-DBCO was prepared. p-NH2-Bn-DTPA (Macrocyclics) was dissolved in 100 μL DMF (0.013 mmol, 8.6 mg), and 87 μL of a 100 mg/mL solution DBCO-PEG4-NHS in DMF was added (0.013 mmol, 8.7 mg). Next, 30 μL triethylamine was added and the reaction was mixed overnight at room temperature. Formation of the product was confirmed using Maldi-ToF with α-Cyano-4-hydroxycinnamic acid as the matrix. The crude reaction mixture was further purified using HPLC with a triethylammonium bicarbonate buffer/methanol solvent system. (0.9 mg, 0.87 μmol, 9%). Maldi-ToF m/z calculated for C51H65N6O17 [M+H]^+^ 1033.441, found 1033.272. Next, DTPA-PEG4-DBCO was dissolved in MilliQ to a 9 mg/mL concentration and added during the coupling of proteins to the IFs when needed, typically 2 eq of DTPA-PEG4-DBCO irt free azides was used. Labeling of the DTPA-bearing IFs with ^111^In was performed using the following protocol: DTPA-IFs were incubated with [^111^In]InCl3 in 0.5 M 2-(*N*-morpholino)ethanesulfonic acid (MES) buffer, pH 5.5 at 37°C, under rotation (550 rpm) for 30 min. PIC were radiolabeled at a ratio of 4 MBq ^111^In to 1 μg of PIC. To complex unbound ^111^In, ethylenediaminetetraacetic acid (EDTA) was added to a final concentration of 5 mM to. The labeling efficiency was determined using instant thin-layer chromatography on silica gels chromatography strips (ITLC-SG; Agilent Technologies) using 0.1 M sodium citrate buffer, pH 6, as mobile phase. Before in vivo administration, radiolabeled IFs were diluted with PBS (pH 7.4) to adjust to a volume of 200 μl/animal.

### Cell lines

The murine melanoma cell line B16-OVA was cultured in RPMI 1640 supplemented with 10% FBS, 2mM L-Glutamine and 0.5% A/A, 1mg/mL Geneticin™ (Gibco, 11811064) and 60 μg/mL Hygromycin B (Gibco, 10687010). Cells were split at a confluency of 70-80% by washing in 1x PBS followed by incubation until detachment in Trypsin-EDTA (BD Biosciences, 215240). Cells were incubated at 37°C, 5% CO2, humidified atmosphere.

### Mice

Mice were housed at the Central Animal Laboratory (Nijmegen, the Netherlands) in accordance with the European legislation. All conducted protocols were approved by the local and national authorities (CCD, The Hague, the Netherlands; license number 10300-2015-0019 and 10300-2019-0020) for the care and use of animals with related codes of practice. Mice were female and between 6 - 10 weeks old at the start of the experiment. Mice were housed in IVC blueline or greenline cages with a maximal number of 6 mice per cage, and were provided with ad libitum food and water and cage enrichment.

Animal studies in Oxford University (for the experiments with IF-A2^(NY-ESO-V)^ and IF-A2^(NY-ESO-V)^/IL-2) were conducted in accordance with the approval of the United Kingdom Home Office. All procedures were done under the authority of the appropriate personal and project licenses issued by the United Kingdom Home Office license number PBA43A2E4. In all experiments, mice were randomized to treatment and outcome assessment was blinded.

### Cell culture of primary murine CD8^+^ T cells

Murine OT-I CD8^+^ T cells or 1G4 CD8^+^ T cells were isolated from the spleen and lymph nodes of OT-I mice (C57BL/6-Tg(TcraTcrb)1100Mjb/Crl (Charles River)) or A2Eso1G4 HHD mice, respectively. Spleens and lymph nodes were digested with 20 ug/mL DNAse I (Roche, 11284932001) and 1 mg/mL collagenase III (Worthington, LS004182) for at least 30 min at 37°C. Organs were meshed through a 100 μm cell strainer and red blood cells were lysed with in-house ammonium-chloride-potassium (ACK) buffer. CD8^+^ T cells were isolated with negative selection using the CD8a^+^ T Cell Isolation Kit, mouse (Miltenyi Biotec, 130-104-075) following manufacturer’s instructions. Cells were counted and diluted to desired concentration and cultured in RPMI 1640 (Gibco, 42401042) with 10% FBS, 2 mM L-Glutamine (Lonza Biowhittaker®, BE17-605E/U1), 0.5% Antibiotic/antimycotic (Gibco, 15240-062) and 50 μM β-mercaptoethanol (Gibco, 21985023) in a 96 well U-bottom plate at 37°C, 5% CO2. Cells used for proliferation studies were stained with 2.5 μM CellTrace Violet (CTV, Invitrogen, C34557) in 1% FBS in 1xPBS for 10 min at 37°C and subsequent recovery for 30 min in 50% FBS at 37°C. Cells were washed in 1xPBS and diluted to desired concentration in cell culture medium. Murine 1G4 CD8^+^ T cells were processed as described above, except in that cells were frozen and thawed before functional assays. Cells were isolated from A2Eso1G4 HHD recipient mice, that were generated as described previously.^[26]^

### Ex vivo CD8^+^ T cell activation assays

IFs were diluted in cell culture medium to desired concentration (see Table 1) and subsequently added to the culture medium. Cells were incubated with IFs until a specific timepoint was reached according to the experimental set-up. Cell supernatant was taken at indicated timepoints and stored at -20°C. Supernatant of murine 1G4 cells was taken on day 2. For wash off experiments, cells were incubated for either 30 min or 90 min with IFs or free pMHC until the cells were transferred under sterile conditions to a 96 well V-bottom plate, washed 2x in 1xPBS and resuspended in cell culture medium. As a control, cells were left untouched until further processing. Supernatant was taken on day 2, replaced with new cell culture medium and proliferation was assessed on day 3.

### Ex vivo cytotoxicity assay

B16-OVA cells were treated with 100 ng/mL murine IFNγ (Peprotech, 315-05) overnight. The next day, cells were harvested and stained with CellTrace™ Violet for some assays. Next, 10.000 B16-OVA cells were added to 96 well U-bottom plates and left to attach for 2-3 hours. CD8^+^ T cells were stimulated for around 20 hours, harvested using PBS with 2mM UltraPure™ 0.5M EDTA, pH 8.0 (Invitrogen, 15575-020) and the desired number was added in 100 μl cell culture medium with 50 μM β-mercaptoethanol to the B16-OVA cells. As a control for background B16-OVA cell death, no T cells were added to the culture. After an additional 24 hours, cells were harvested by taking off the medium, washing in 1xPBS and incubating with 30 μl Trypsin-EDTA until all cells were detached. Cells were processed for flow cytometry. The percentage lysis was calculated using the following formula: % lysis = (1-(Freq. treated viable target cells/Freq. no T cell viable target cells))x100.

### Antibodies and Reagents in flow cytometry

For flow cytometric analysis, cells were stained for 20-30 min at 4°C in the dark using 1:2000 eFluor 780 fixable viability dye (eBiosciences, 65-0865-014), Propidium Iodide (Miltenyi Biotec, 130-093-233) or 1:2000 7-AAD (eBiosciences, 00-6993-50). Next, cells were stained for 20-30 min with antibody mixtures. Murine OT-I CD8^+^ T cells of ex vivo assays were stained with: aCD8a-PE (BD Biosciences, 553033), aCD8a-APC (Biolegend, 100712), aCD25-APC (eBiosciences, 17-0251-82), aCD69-BV510 (Biolegend, 104531). The 1G4 murine CD8^+^ T cells were additionally stained with: mouse-anti-human TCR Vβ 13.1-PE (clone: H131, Biolegend, 362409). For in vivo studies, cells isolated from selected organs were stained with aCD11b-PE (Biolegend, 101208), aCD45.1-PerCpCy5.5 (Biolegend, 110726), CD3-APC-Cy7 (Biolegend, 100221), aCD8-PE-Cy7 (Biolegend, 100722) and αCD16-αCD32 FcR block (BD Biosciences, 553142). Next, cells were washed 2x in PBA and transferred to polystyrene flow cytometry tubes. In cases were 7-AAD or PI was used, samples were stained 10 min at 4°C before run in 1xPBS. Human transfected HLA-A2.1^+^ CD8^+^ T cells were stained with: anti-mouse TCR-β-FITC (Biolegend, 109205), anti-mouse TCR-β-BV421 (Biolegend, 109230), aCD25-PE-Cy7 (Biolegend, 302612), aCD8-BV510 (Biolegend, 344732). Samples were run on the BD FACSVerse (BD Biosciences) and analyzed using FlowJo vX0.7 after compensation using AbC Total Antibody Compensation Bead Kit (Invitrogen, A10497). Cell count was determined using Precision Count Beads (Biolegend, 424902). The division index was determined by manually gating on CTV peaks and using the following formula: 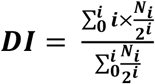 (where i is the division cycle and N is the proportion of cells in this division cycle.)

### Cytokine levels in supernatant

Cytokine levels were measured in cell supernatant using an enzyme linked immunosorbent assay (ELISA). Murine IFNγ levels and IL-2 levels were measured using the Mouse Interferon gamma (IFNγ) Uncoated ELISA Kit (Thermo Fisher Scientific, 88-7314-76) or IL-2 Mouse Uncoated ELISA Kit (Thermo Fisher Scientific, 88-7024-76) following the manufacturer’s instructions. The supernatants of human CD19 CAR T cells or human TCR transfected CD8^+^ T cells were analyzed by IFNγ Human Uncoated Elisa Kit (Invitrogen, Thermo Fisher Scientific # 88-7316-88). Plates were read on BioRad plate reader at a wavelength 450 nm and subtracted by 595 nm.

### Generation and culture of Flt3 ligand dendritic cells for RNAseq

Legs of C57BL/6 mice (Charles River) were collected. Bone marrow was washed out onto a 100 μm cell strainer, cells were washed and plated in 10 cm petri dish (Greiner, 633185) in supplemented cell culture medium was described above. The RPMI medium was additionally supplemented with 200 ng/mL human Flt3 ligand (Miltenyi Biotec, 130-096-479) and 5 ng/mL murine GM-CSF (Peprotech, 315-02) and 50 μM β-mercaptoethanol. Cells were cultured at a density of 15×10^6^ per plate at 10% CO2, 37°C in humidified atmosphere. After 5 days, medium was replaced with fresh 200 ng/mL Flt3 ligand and 5 ng/mL murine GM-CSF. After 9 days, cells were harvested and replated at a density of 3×10^6^ cells per plate, and 200 ng/mL fresh Flt3 ligand and 5 ng/ml GM-CSF containing medium was added up to 10 ml. One day before T cell co-culture, cells were stimulated with 0.3 μg/ml LPS (Invivogen, vac-3pelps). On the day of T cell co-culture, cells were given 100 ng/ml SIITFEKL peptide (Genscript) for 3 hours.

### Bulk RNAseq experiment

For RNA sequencing experiments, OT-I CD8^+^ cells were incubated for 8 hours and 22 hours with 5 ng IF-pMHC^(SIIT)^/IL-2 or SIITFEKL-pulsed Flt3L DCs in a ratio of five OT-I CD8^+^ cells to two DCs in 37°C, 5% CO2, humidified atmosphere. Viable cells were sorted on the BD FACSMelody into polypropylene tubes based on CD8-APC (Biolegend, 100721), CD103-PE (Biolegend, 121405) expression. Cells were lysed in 200 μl TRIzol (Thermo Fisher, 15596026) and stored at -80°C before sending to Single Cell Discoveries B.V. (Utrecht, the Netherlands) for RNA extraction and bulk RNA sequencing. All samples passed the quality control with high quantity and quality, as assessed by Agilent Bioanalyzer. Sequencing was performed using an adapted version of the CEL-seq protocol. In brief: Total RNA was extracted using the standard TRIzol protocol and used for library preparation and sequencing. mRNA was processed as described previously, following an adapted version of the single-cell mRNA seq protocol of CEL-Seq.^[42,43]^ In brief, samples were barcoded with CEL-seq primers during a reverse transcription and pooled after second strand synthesis. The resulting cDNA was amplified with an overnight in vitro transcription reaction. From this amplified RNA, sequencing libraries were prepared with Illumina Truseq small RNA primers. The DNA library was paired-end sequenced on an Illumina Nextseq™ 500, high output, with a 1×75 bp Illumina kit (R1: 26 cycles, index read: 6 cycles, R2: 60 cycles). For data analysis, Read 1 was used to identify the Illumina library index and CEL-Seq sample barcode. Read 2 was aligned to the Mouse mm10 + mitochondrial genes reference transcriptome using BWA MEM.^[44]^ Reads that mapped equally well to multiple locations were discarded. Mapping and generation of count tables was done using the MapAndGo script1. A complete overview of the genes significantly different between DCs and IF-pMHC^(SIIT)^/IL-2 is provided in **Table S1**.

### Transfection of human T cells with TCR

CD8^+^ T cells obtained using Ficoll density gradient centrifugation (Lymphoprep, ELITechGroup) followed by CD8^+^ T cell isolation (CD8^+^ T Cell Isolation Kit, human, Miltenyi Biotec) on buffy coats from HLA-A2.1^+^ human donors after written informed consent in accordance with the Declaration of Helsinki and in agreement with institutional guidelines. CD8^+^ T cells were transfected as previously described^[45]^ with mRNA (5 mg/mL, BioNTech RNA Pharmaceuticals) encoding for a murinized T cell receptor (specific for the HLA-A2.1-specific NY-ESO-1 epitope SLLWITQC). Transfection efficiency was determined 1 day after transfection using anti-mouse TCR-β-FITC (BioLegend, 109205) and was typically between 80 and 90%. Cells were stained with CTV as described above and treated with IF. Supernatant was taken 3 days after stimulation and proliferation was evaluated 4 days after stimulation. For activation studies, cells were not stained with CTV and activation was read out one day after IF stimulation. Cells were cultured in X-Vivo 15 (BE-02-060F, Lonza) with 2% human serum.

### Stimulation of CD19 CAR T cells

Human CD3^+^ T cells were obtained from buffy coats of healthy donors by negative selection using RosetteSep Human T Cell Enrichment Cocktail Kit (STEMCELL Technologies) and stored at -80ºC until use. T cells were cultured with X-VIVO 15 Cell Medium (Cultek, BE02-060Q) supplemented with 5% AB human serum (Sigma, H4522), penicillin-streptomycin (100 mg/mL), and IL-2 (50 IU/mL; Miltenyi Biotec), and stimulated with Dynabeads (Gibco, 11131D). After 24 hours of T cell stimulation with beads, cells were transduced with anti-CD19 CAR lentiviral particles at a multiplicity of infection (MOI) of 5 and cultured for 6 days. Dynabeads were removed and supplemented medium was added to cells. T cell culture was maintained with X-vivo + 5% AB human serum + 100 mg/mL penicillin-streptomycin for a period of 48h. Then, the expression of anti-CD19 CAR was assessed by cell-based fluorescence using Allophycocyanin (APC) AffiniPure F(ab’)2 Fragment Goat Anti-Mouse IgG, F(ab’)2 Fragment Specific (Jackson Immunoresearch #115-136-072) before adding IF. The expression of anti-CD19 CAR on T cells in all experiments were ranging between 40-50%. Afterwards, IFs were added to CD19 CAR T cell culture (1×10^6^ cells/ml) at a final concentration of 2 μg/mL. CD19 CAR T cells were split during the stimulation with IFs and supplemented X-VIVO without IL-2. The supernatant of the CD19 CAR T cells was collected at different timepoints (24 hours, 72 hours and 5 days after adding IF).

### In vivo biodistribution and SPECT/CT imaging

C57BL/6 mice received 290 ng pMHC intravenously either bound or unbound to IFs or blank IFs labelled with Indium-111. After 15 min, 1 hour, 2 hours, 4 hours, 8 hours, 24 hours, 48 hours and 72 hours blood was collected from part of the mice through thigh bone puncture. After 24 hours or 72 hours, mice were sacrificed by CO2 asphyxiation and some mice were used for SPECT-CT analysis as described below. Organs were harvested from mice and radioactivity was measured using gamma counter (Wizard, PerkinElmer). The % of injected dose per gram tissue/organs was calculated from the amount of radioactivity measured in aliquots of the injected dose. The %ID of lymph nodes was calculated using the similar formular but not normalized to weight. The %ID muscle was adjusted to the average weight of lymph nodes which was calculated to be 3.5 mg. At 24 hours and 72 hours after injection of ^111^In labeled IFs, one mouse of each group was imaged with SPECT/CT after euthanization. Images were acquired for 2 hours with the U-SPECT-II/CT (MILabs, Utrecht, The Netherlands) using a 1.0 mm diameter pinhole mouse high sensitivity collimator, followed by CT scan (615 μA, 65 kV) for anatomical reference. Scans were reconstructed with MILabs reconstruction software using a 16-subset expectation maximization algorithm, with a voxel size of 0.4 mm and 1 iteration. SPECT/CT scans were analyzed and maximum intensity projection (MIP) were created using Inveon Research Workplace software (Siemens).

### In vivo CD8^+^ T cell proliferation

For proliferation studies with OT-I T cells, CD8^+^ OT-I T cells were isolated and labelled with CTV according to the aforementioned protocol. Depending on the experimental set-up, 0.5×10^6^ or 1×10^6^ CTV labelled cells were injected intravenously. One day later, the amount as indicated in the figure legend was injected either iv or sc. After 3 or 4 days, spleen and lymph nodes were isolated and processed for flow cytometry. Samples were run on the MACS Quant or BD FACSVerse and gated for live cells.

For proliferation studies with 1G4 T cells, 1G4 T cells were isolated using “untouched isolation” MACS microbead selection (Miltenyi Biotec). Cells were labeled labelled with CTV by mixing 1:1 in PBS, cells at 2×107 /mL with CTV at 5 μg/mL (Thermo Fisher) and incubating for 15 min at 37 °C followed by blocking in FBS and washing with complete medium according to the aforementioned protocol. Depending on the experimental set-up, 1×10^6^ CTV labelled cells were injected intravenously. One day later, the amount as indicated in the figure legend was injected intravenously. After 3 or 4 days, spleen and lymph nodes were isolated and processed for flow cytometry. Samples were run on the MACS Quant, BD FACSVerse or BD LSR Fortessa and gated for live cells.

### Lung metastasis model

C57BL/6 mice were injected iv with 0.25 ×10^6^ freshly isolated OT-I CD8^+^ T cells or 1x PBS. The next day, mice received 1.4 μg pMHC either bound to IFs or free pMHC or PBS iv After 4 days, 0.8×10^6^ B16-OVA cells, pre-treated over night with 100 ng/ml murine IFNγ, as described above, were added in 1x PBS iv After 14 days post tumor cell injection, mice were euthanized using CO2 asphyxiation, and lungs were perfused with cold PBS 2mM EDTA through the right ventricle of the heart. The lungs were fixed in in-house Fekete’s solution and lung metastases were counted the next day under blinded conditions. Metastatic burden on the lung surface was quantified by dividing the total metastatic area by total lung area using ImageJ Fiji software.^[46]^

### Subcutaneous tumor model

C57BL/6 mice were injected sc with 100 μl of 0.25×106 B16-OVA cells in matrigel. When tumors had reached a size between 50-100 mm3 mice were randomized according to tumor volume and 0.4×106 OT-I T cells were adoptively transferred intravenously. One day later, 200 μg InVivoMab rat-anti-mouse PD-1 antibody (clone: RMP1-14, BioXCell, BE0146) was injected in InVivoPure pH 7.0 Dilution Buffer (BioXCell, IP0070) intraperitoneal in some mice, which was given again on day 4 and 7. One day after OT-I T cell transfer, mice either received 0.5 μg of pMHC(SIIN) on IFs or blank IFs in similar amounts as to the IF-pMHC(SIIN), diluted in sterile PBS sc around the tumor. Tumor sizes were measured every other day with a caliper. Width x length x depth x 0.4 was used to calculate tumor volumes. Tumor growth was followed up and mice were sacrificed when the tumor reached ≥1500 mm^3^ threshold. For the representation of tumor growth graphs, tumor volumes of dead mice were kept at the threshold value after euthanization until the end of the experiment. Tumors were formalin-fixed (10% formalin) and paraffin-embedded (FFPE) for histological analysis.

### Immunohistochemistry

Slides of 4 μm of FFPE B16-OVA tumors were cut. Next, slides were deparaffinized and antigens were retrieved using ENVISION Flex Target Retrieval solution (Dako Omnis, K8004) in a microwave (3 min 1000W, 20 min 180W). Endogenous peroxidase was blocked 10 min in 3% H2O2 (Merck, 107209) washed and then blocked in 1% bovine serum albumin in TBS-T for 10 min. Afterwards, slides were incubated with rabbit-anti-mouse CD8a (dilution: 1/1000, clone: EPR21769, Abcam, ab217344) for 1 hour at room temperature. Secondary BrightVision® poly-HRP-anti-rabbit antibody (dilution: 1/2, ImmunoLogic, S/DPVR-HRP) was applied for 30 min at room temperature. Next, Envision Flex HRP Magenta Substrate Chromogene System (Dako Omnis, DM857) was applied for 5 minutes. Reaction was stopped by washing for 1 min in tap water. Between each step, slides were rinsed in TBS-T. Slides were counterstained in hematoxylin and enclosed with QuickD mounting solution (Klinipath, 7281).

### Tissue Imaging and quantitative digital analysis

Whole tissue slides were imaged with the Vectra Intelligent Slide Analysis System (Version 3.0.4, PerkinElmer Inc.) as previously described^[47]^. Phenochart (Version 1.1.0, Akoya Biosciences) was used to select the tumor area for analysis. Spectral libraries were built based on unstained tissue, melanin pigmentation and single staining consisting of hematoxylin for nuclear staining and EnvisionFlex magenta for CD8^+^ T cells. Training of the inForm Advanced Image Analysis Software (Version 2.4.8, Akoya Biosciences) was performed on a selection of 10 to 15 representative original multispectral images to discriminate between tumor and necrotic areas, cell segmentation and phenotyping of CD8^+^ T cells, melanin-pigmented cells and other cells (**Figure S10**). Batch analysis of multiple original multispectral images of the same tumor was allowed by saving settings applied to the training images within an algorithm. The numbers of intratumoral CD8^+^ T cells were quantified and normalized for the tumor area (cells/mm^2^).

### Statistics

All data is represented as mean ± standard error of the mean (SEM). Graphs were generated in GraphPad Prism (Version 8.0.2) or R Studios (Version 4.0.3). Statistical analysis was performed on transformed data where appropriate, using Graph Pad Prism with the appropriate testing methods as indicated in the figure legends. Statistical significance was defined as a two-sided significance level of <0.05. Only p-values <0.05 are indicated in the graphs.

## Supporting Information

Supporting Information is available from the Wiley Online Library or from the author.

## Acknowledgements

L.W. and J.W. contributed equally to this work. C.G.F. and R.H. contributed equally to this work. We thank the central animal laboratory Nijmegen (CDL) for their contribution to the animal work. We would like to acknowledge Mireille Toebes and Ton N. Schumacher for providing the H-2kb, HLA-A2 and hβ2m constructs and protocols, and we want to acknowledge Ugur Sahin and Mustafa Diken for providing the murinized T cell receptor specific for the HLA-A2.1-specific NY-ESO-1 epitope SLLWITQC.

## Data availability statement

The RNA sequencing raw data is under submission for the GEO database. An overview of the normalized counts of all detected genes and of significantly different genes after 8 hours and 22 hours can be found in Table S1.The rest of the data is available from the authors upon reasonable request.

## Ethics approval statement

Mice were housed at the Central Animal Laboratory (Nijmegen, the Netherlands) in accordance with the European legislation. All conducted protocols were approved by the local and national authorities (CCD, The Hague, the Netherlands; license number 10300-2015-0019 and 10300-2019-0020)for the care and use of animals with related codes of practice. Animal studies in Oxford University were conducted in accordance with the approval of the United Kingdom Home Office. All procedures were done under the authority of the appropriate personal and project licenses issued by the United Kingdom Home Office license number PBA43A2E4.

## Patient consent statement

All subjects gave written informed consent in accordance with the Declaration of Helsinki and in agreement with institutional guidelines.

