## Supporting Information for "Immunofilaments Provide a Nanoscale Platform for In Vivo T Cell Expansion and Cancer Immunotherapy"

**Figure S1.** Multivalent immunofilaments presenting pMHC$^{(SIIN)}$ support OT-I T cell activation, proliferation and target cell killing.
**Figure S2.** Immunofilaments presenting pMHC$^{(SIIT)}$ and in particular IF-pMHC$^{(SIIT)}$/IL-2 support OT-I T cell viability and proliferation
**Figure S3.** Characterization of OT-1 and comparison of differential genes of OT-I T cells after 22 hours stimulation with DCs and IF-pMHC$^{(SIIN)}$/IL-2.
**Figure S4.** Immunofilaments presenting A2$^{(NY\text{-}ESO\text{-}V)}$/IL-2 stimulate TCR-transfected human HLA-A2.1$^{+}$ CD8$^{+}$ T cells in a specific manner.
**Figure S5.** Impact of single versus co-presentation of ProL, αCD28 and IL-2 on IFs for CD19 CAR T cell activation.
**Figure S6.** The in vivo biodistribution and T cell activation of IF-pMHC$^{(SIIN)}$ in WT mice adoptively transferred with WT CD8$^{+}$ T cells or OT-I CD8$^{+}$ T cells.
**Figure S7.** Immunofilaments presenting pMHC$^{(SIIN)}$ expand antigen-specific OT-I T cells in vivo.
**Figure S8.** Immunofilaments presenting A2$^{(NY\text{-}ESO\text{-}V)}$ expand antigen-specific 1G4 T cells adoptively transferred into recipient mice in vivo.
**Figure S9.** Tumor growth and CD8$^{+}$ T cell infiltration in sc B16-OVA tumors.
**Figure S10**. Representative example of image analysis approach of T cell infiltration into B16-OVA tumors.
**Table S1**. Normalized counts by RNAseq for all detected genes and of significantly different genes between DCs and IF-pMHC$^{(SIIT)}$/IL-2 after 8 hours and 22 hours stimulation of OT-I T cells.

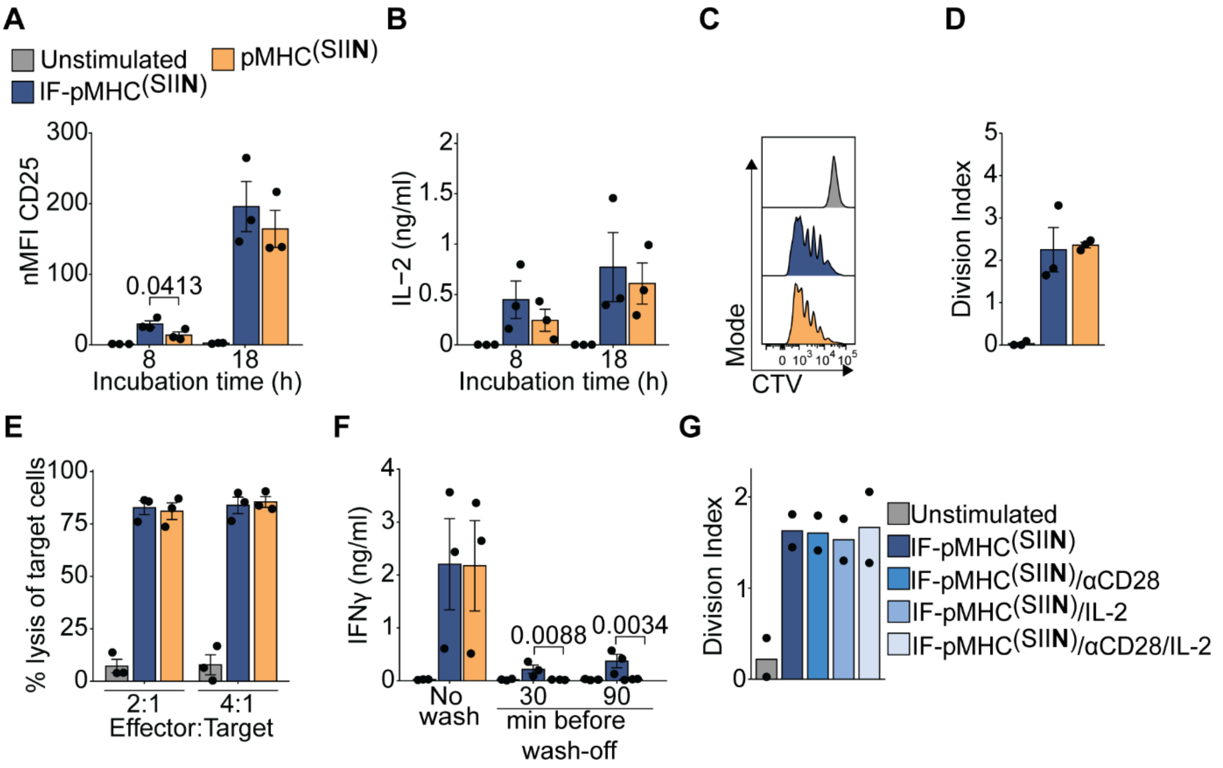


**Figure S1. Multivalent immunofilaments presenting pMHC$^{(SIIN)}$ support OT-I T cell activation, proliferation and target cell killing.** (A) Flow cytometry quantification of the normalized geometric mean fluorescence intensity (nMFI) of activation marker CD25 after 8 hours or 18 hours. Statistical significance was determined by two-way ANOVA on log-transformed data with post-hoc Sidak's multiple comparison test. (B) IL-2 production by OT-I T cells after 8 hours and 18 hours, determined by ELISA. Statistical significance was tested by two-way ANOVA on log-transformed data. (C,D) Proliferation by CTV dilution (C) and quantification of division index (D) of non-stimulated OT-I T cells or stimulated for 3 days with IF-pMHC$^{(SIIN)}$ or pMHC$^{(SIIN)}$. Statistical significance was tested with unpaired t-test. (E) Flow cytometry quantification of the percentage of lysed B16-OVA melanoma target cells 24 hours after co-incubation with OT-1 T cells pre-stimulated for 20 hours. Statistical significance was tested with two-way ANOVA on log-transformed data. (F) IFNγ production by OT-I T cells after two days by ELISA, either without washing away the stimulation (left) or after washing the cells either 30 min or 90 min after addition of IF-pMHC$^{(SIIN)}$ or pMHC$^{(SIIN)}$. Statistical significance was tested with two-way ANOVA on log-transformed data and post-hoc Sidak's multiple comparison test. (A-F) $n$ = 3 in three independent experiments. p-values are indicated in the Figure. (G) Flow cytometry quantification of the division index of OT-I T cells stimulated for 3 days comparing various IF-pMHC$^{(SIIN)}$. $n$ = 2 in two independent experiments.

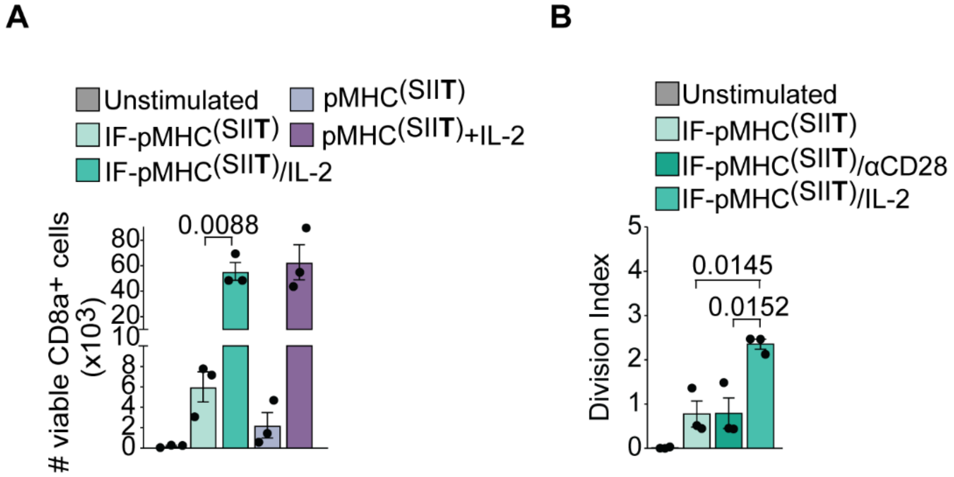


**Figure S2. Immunofilaments presenting pMHC$^{(SIIT)}$ and in particular IF-pMHC$^{(SIIT)}$/IL-2 support OT-I T cell viability and proliferation.** (A) Quantification of the number of viable cells by flow cytometry after 3 days of stimulation with various IF-pMHC$^{(SIIT)}$. Statistical significance was determined by one-way ANOVA and post-hoc Tukey's multiple comparison test. (B) Flow cytometry quantification of the division index of OT-I T cells stimulated for three days comparing various IF-pMHC$^{(SIIT)}$. Statistical significance was determined by one-way ANOVA and post-hoc Tukey's multiple comparison test. (A-B) $n$ = 3 in three independent experiments. p-values are indicated in the Figure.

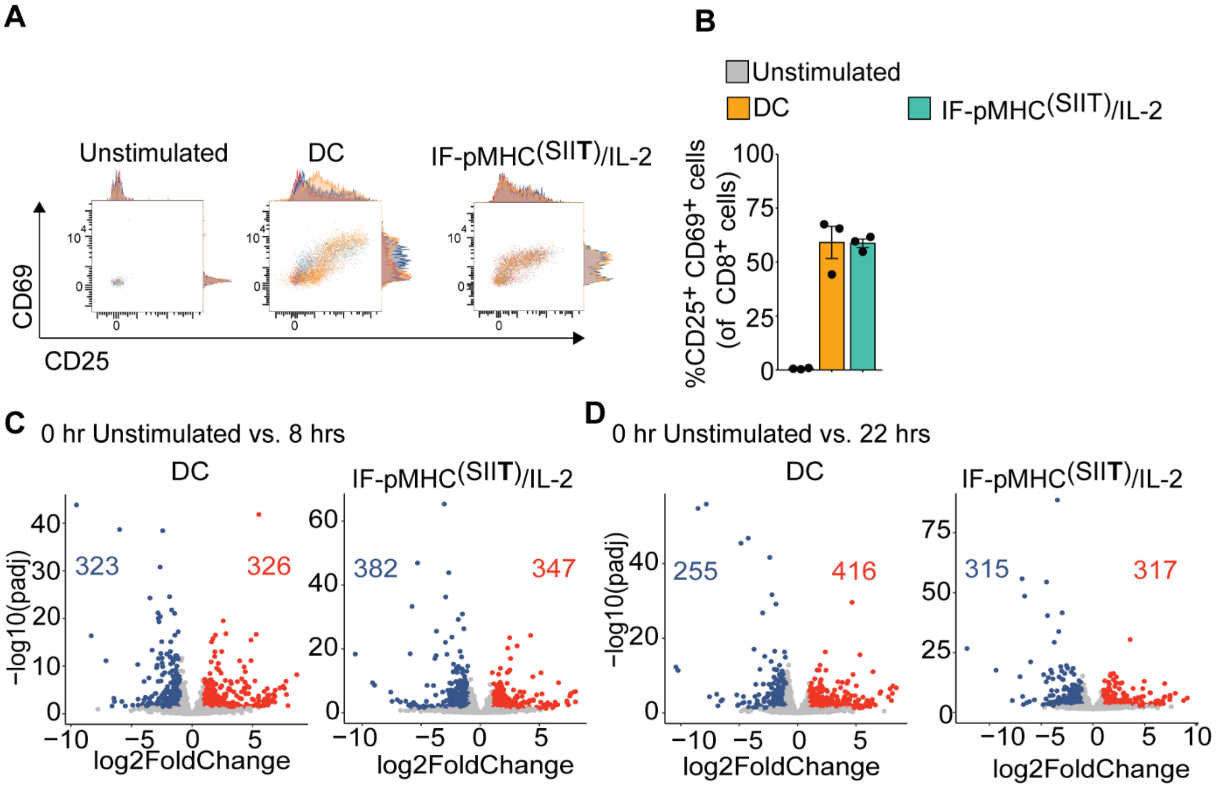


**Figure S3. Characterization of OT-1 and comparison of differential genes of OT-I T cells after 22 hours stimulation with DCs and IF-pMHC$^{(SIIN)}$/IL-2.** (A-B) Representative flow cytometry plots (orange, red and blue represent the three donors) (A) and quantification (B) of the % of CD25$^{+}$/CD69$^{+}$ OT-I T cells before and after 22 hours stimulation with Flt3 ligands DCs and pMHC$^{(SIIN)}$/IL-2. (C-D) Volcano plots depicting differential genes comparing OT-I T cells stimulated by DCs or IF-pMHC$^{(SIIN)}$/IL-2 at 0 hours vs 8 hours (C) and at 0 hours vs 22 hours (D). Genes with corrected $p<0.05$ are highlighted in blue/red.

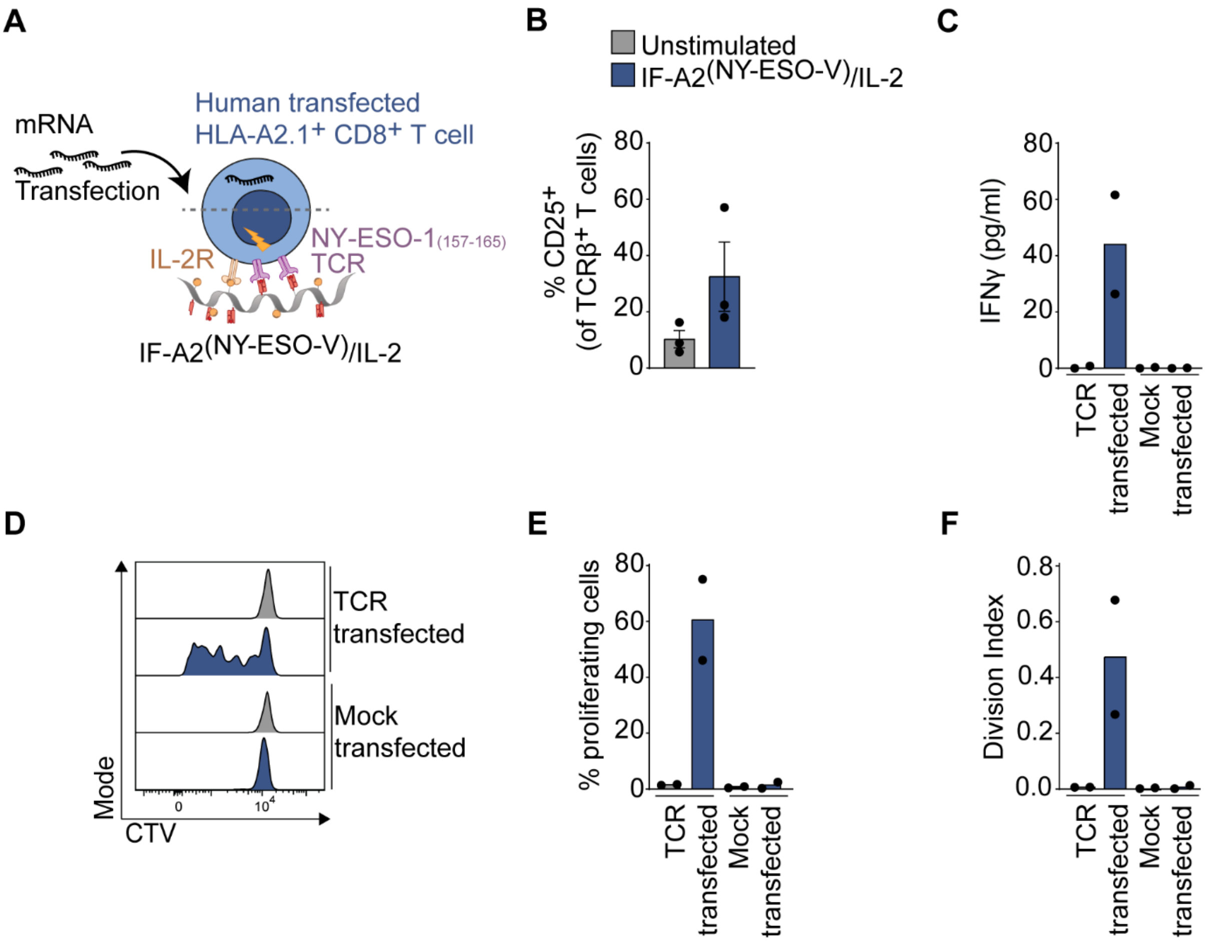


**Figure S4. Immunofilaments presenting A2$^{(NY\text{-}ESO\text{-}V)}$/IL-2 stimulate TCR-transfected human HLA-A2.1$^{+}$ CD8$^{+}$ T cells in a specific manner.** (A) Schematic overview of stimulation of human HLA-A2.1$^{+}$ CD8$^{+}$ T cells transfected with TCR specific for NY-ESO-1$_{(157\text{-}165)}$ with IF-A2$^{(NY\text{-}ESO\text{-}V)}$/IL-2. (B) Flow cytometry quantification of the percentage of activated CD25$^{+}$ T cells of all CD8$^{+}$ transfected T cells after 24 hours. (C) IFNγ production after 3 days of incubation with IF. *n* = 3 in three independent experiments. (D-F) Proliferation by CTV dilution (D) and quantification of the percentage of proliferating cells (E) and division index (F) TCR-transfected or mock-transfected T cells left non-stimulated or stimulated for 4 days with IF-A2$^{(NY\text{-}ESO\text{-}V)}$/IL-2. (C, E, F) *n* = 2 in two independent experiments.

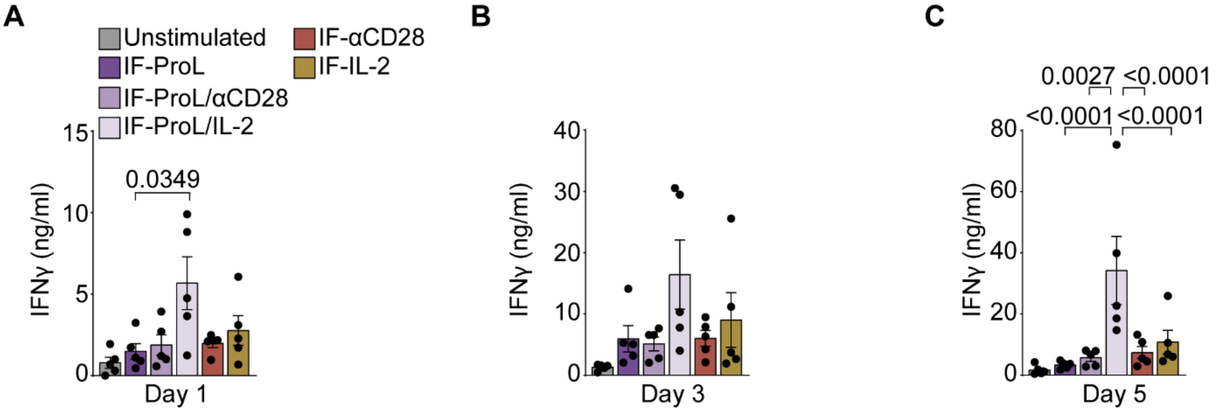


**Figure S5. Impact of single versus co-presentation of ProL, αCD28 and IL-2 on IFs for CD19 CAR T cell activation.** (A-C) IFNγ production after 1 (A) , 3 (B) and 5 days (C) incubation with IF. Statistical significance was determined by two-way ANOVA on log-transformed data with post-hoc Tukey's multiple comparison test. $n = 3$ in three independent experiments. p-values are indicated in the Figure.

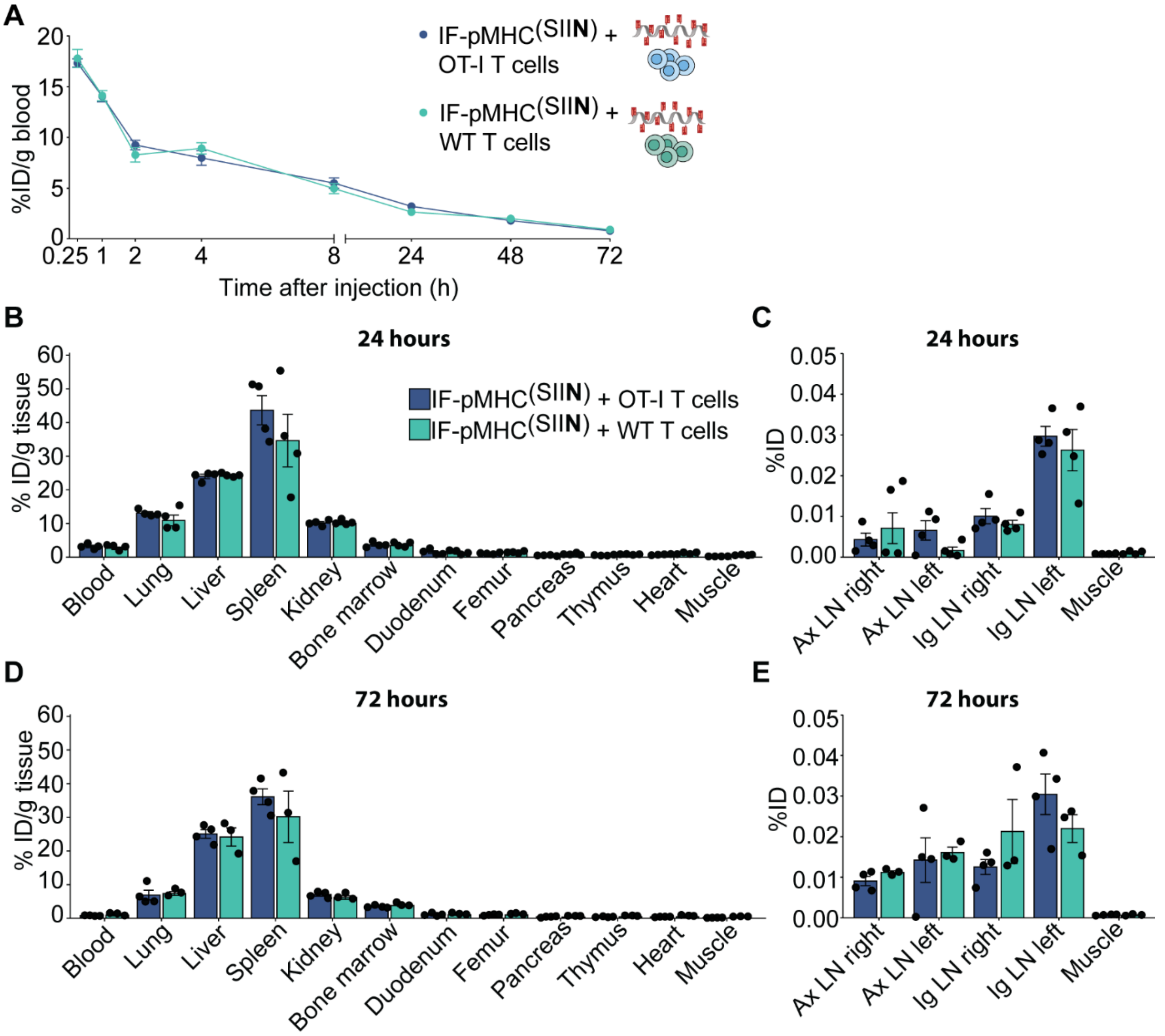


**Figure S6. The in vivo biodistribution and T cell activation of IF-pMHC$^{(SIIN)}$ in WT mice adoptively transferred with WT CD8$^{+}$ T cells or OT-I CD8$^{+}$ T cells.** (A) Kinetics of $^{111}$In-labelled IF-pMHC$^{(SIIN)}$ in blood after iv injection in WT CD45.1$^{+}$ C57Bl/6J mice, expressed at % of injected dose (ID) per gram of blood. Mice received CD45.2$^{+}$ wildtype T cells or CD45.2$^{+}$ OT-I T cells one day prior to IFs administration. Statistical significance was determined with mixed effect analysis and post-hoc Sidak's multiple comparison test. $n$ = 5 for t=0.25,1,2,4, 8 and 24 hours in one experiment. (B-E) Quantitative ex vivo analysis of the biodistribution of IF-pMHC$^{(SIIN)}$ 24 hours (B-C) and 72 hours (D-E) after iv injection across different organs expressed at % of ID per gram of tissue. Statistical significance was determined per timepoint by two-way ANOVA on log-transformed data with post-hoc Sidak's multiple comparison test.

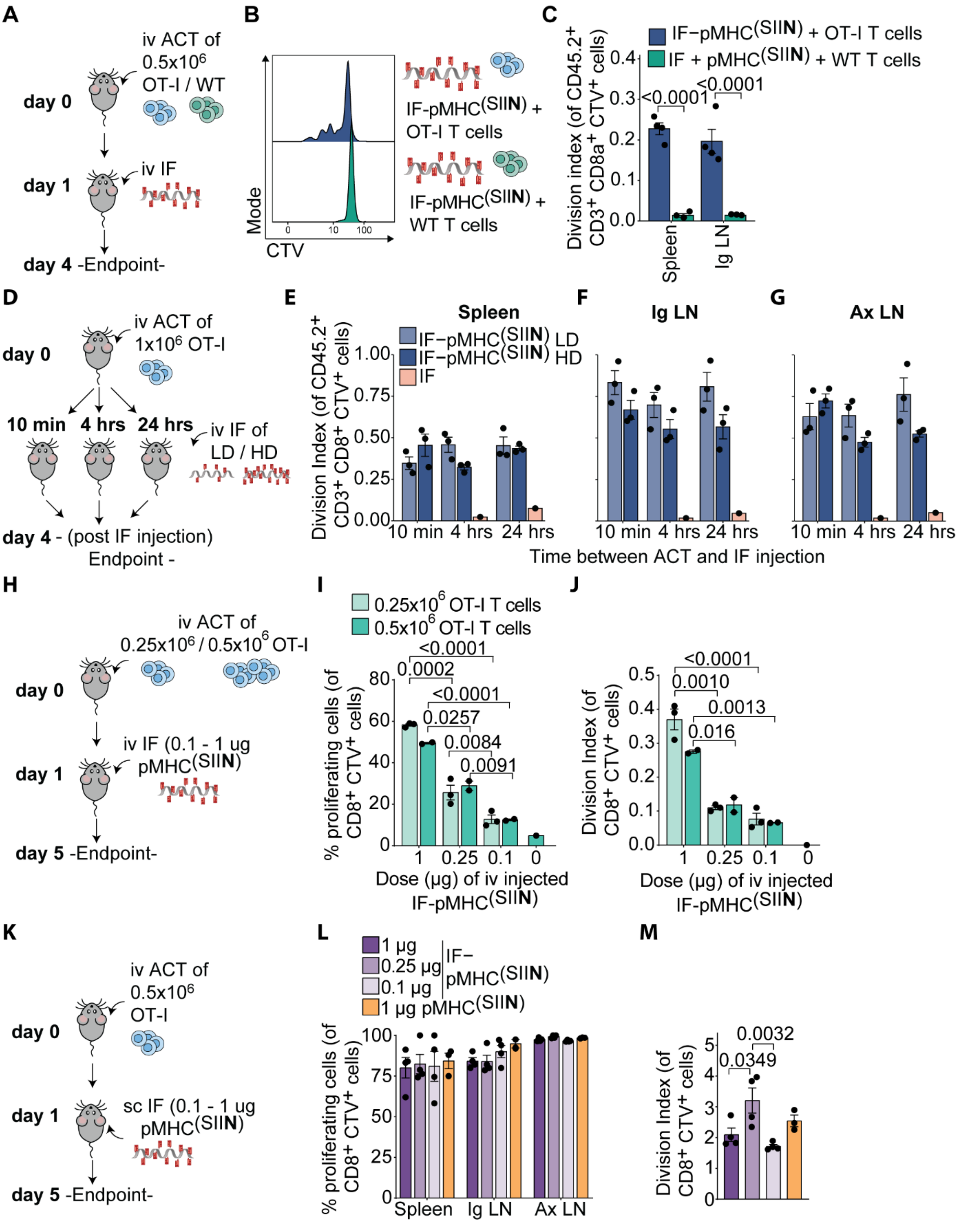


**Figure S7. Immunofilaments presenting pMHC$^{(SIIN)}$ expand antigen-specific OT-I T cells in vivo.** (A) Schematic overview of experiment to determine in vivo OT-I T cell proliferation after iv administration of IF-pMHC$^{(SIIN)}$ or IFs + free pMHC$^{(SIIN)}$. (B-C) Ex vivo flow cytometric analysis of CTV dilution (B) and quantification (C) of the division index of OT-I T cell in the spleen 3 days after iv administration of IF-pMHC$^{(SIIN)}$ (0.29 µg pMHC$^{(SIIN)}$ ) in WT C57Bl/6J adoptively transferred with 0.5x10$^{6}$ CTV-labelled OT-I T cells or WT T cells. Significance was determined on log-transformed data with two-way ANVOA and post-hoc Dunnet's multiple comparison test. $n$ = 3 in one independent experiment. p-values are indicated in the Figure. (D) Schematic overview of experiment to study the effect of time of administration of IFs following OT-I adoptive transfer. (E,F) Quantification of the division index of OT-I T cells in the spleen (E), in the inguinal (Ig) LN and the axillary (Ax) LN (F) after iv administration of IFs alone of IF-pMHC$^{(SIIN)}$ with a low density (LD): ~5 pMHC per IF) or high density (HD): ~16 pMHC per IF). One µg of pMHC$^{(SIIN)}$ on IFs were injected either 10 min, 4 hours or 24 hours after adoptive transfer of OT-I T cells. $n$ = 3 in one experiment. Significance was tested per organ with two-way ANOVA on log-transformed data with post-hoc Sidak's multiple comparison test. (H) Schematic overview of experiment to study impact of OT-I cell number and IF-pMHC$^{(SIIN)}$ dose on OT-I proliferation after iv administration of IF. $n$ = 3 for 0.25x10$^{6}$ OT-I and $n$ = 2 for 0.5x10$^{6}$ OT-1 in one experiment. (I-J) Quantification of the % of proliferating cells (I) and the division index (J) of OT-I T cells (0.25 or 0.5x10$^{6}$) in the spleen after iv administration of IF-pMHC$^{(SIIN)}$ at different doses (dose refers to pMHC$^{(SIIN)}$ amount). Significance was determined by two-way ANOVA on logit-transformed (I) or log-transformed (J) data with post-hoc Tukey's multiple comparison test. p-values are indicated in the Figure. (K) Schematic overview of experiment to study the impact of IF-pMHC$^{(SIIN)}$ dose on OT-I proliferation after sc administration of IF. (L-M) Quantification of the % of proliferating cells (L) in the spleen, Ig LN and Ax LN and the division index (M) in the Ax LN of adoptively transferred OT-I T cells (0.5x10$^{6}$) after sc administration of IF-pMHC$^{(SIIN)}$ at different doses or after sc administration of free pMHC$^{(SIIN)}$. Significance was determined with one-way ANOVA on logit-transformed (L) or log-transformed (K) data and post-hoc Tukey's multiple comparison test. $n$ = 4 in one experiment. p-values are indicated in the Figure.

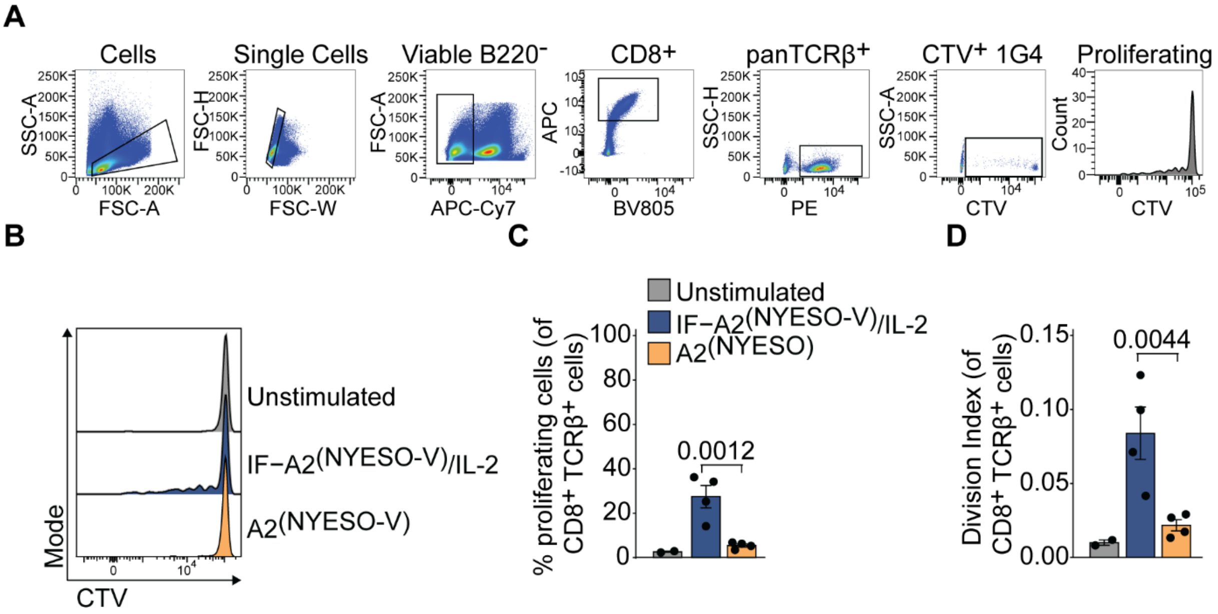


**Figure S8. Immunofilaments presenting A2$^{(NY\text{-}ESO\text{-}V)}$ expand antigen-specific 1G4 T cells adoptively transferred into recipient mice in vivo**. (A) Flow cytometry gating strategy to identify proliferating 1G4 T cells. (B-D) Ex vivo flow cytometric analysis of CTV dilution (B) and quantification of the % of proliferating cells (C) and division index (D) of 1G4 T cells in the spleen 3 days after iv administration of IF-A2$^{(NY\text{-}ESO\text{-}V)/IL2}$ or free A2$^{(NY\text{-}ESO\text{-}V)}$ (1.4 µg A2$^{(NY\text{-}ESO\text{-}V)}$). Statistical significance was determined by unpaired t-test on logit-transformed (C) or log-transformed (D) data. *n* = 4 in one experiment. p-values are indicated in the Figure.

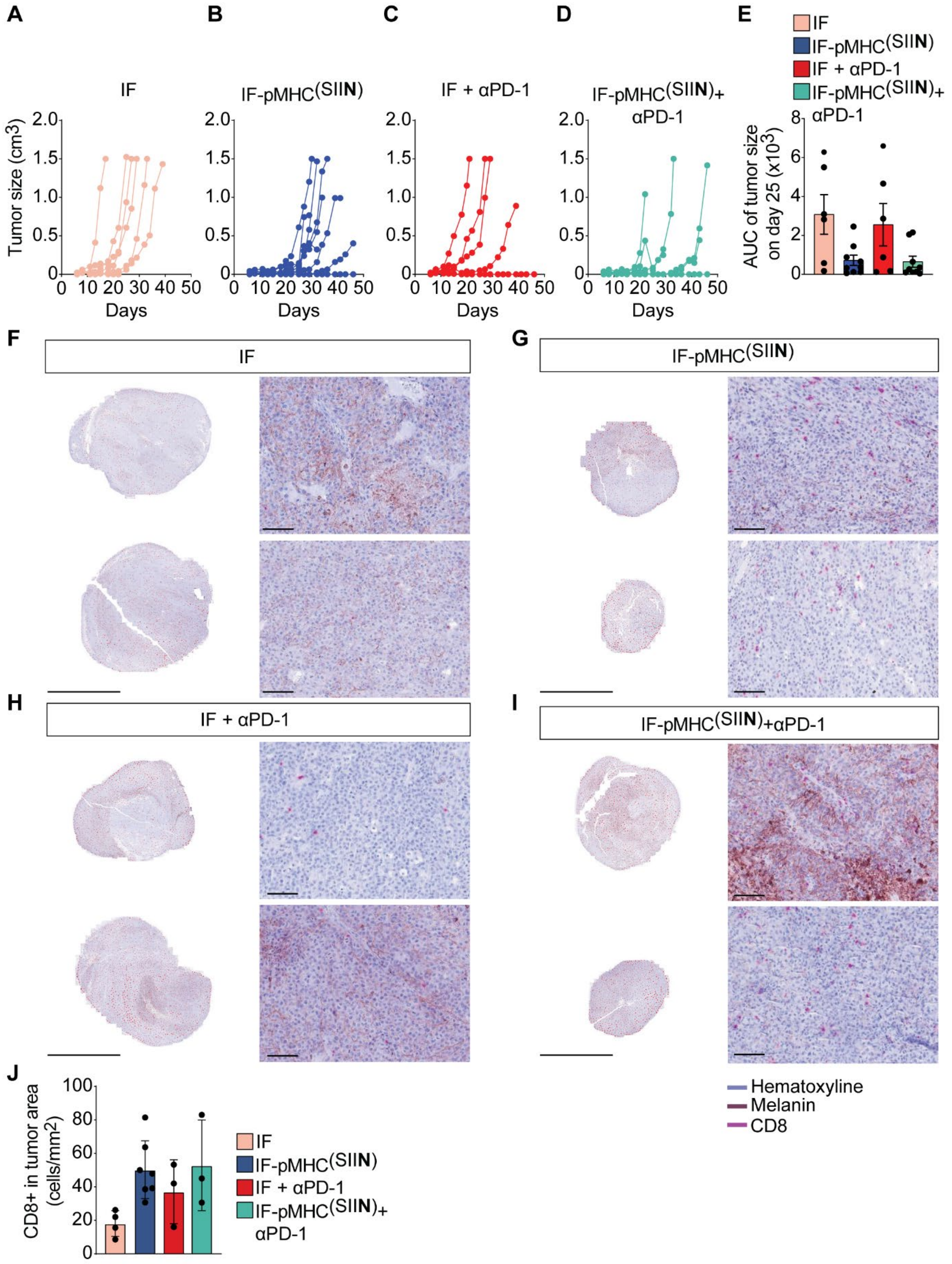


**Figure S9. Tumor growth and CD8$^{+}$ T cell infiltration in sc B16-OVA tumors.** (A-D) Individual curves of quantification of the size of B16-OVA tumors in mice treated sc with non-functionalized IFs (A), IF-pMHC$^{(SIIN)}$ (B), non-functionalized IFs + αPD-1 (C), and IF-pMHC$^{(SIIN)}$ + αPD-1 (D). $n$ = 6-9 in one independent experiment. (E) Quantification of the area under the curve (AUC) of the tumors in the sc model on day 25. Statistical significance was tested by Kruskal-Wallis test on log-transformed data. (F-I) Representative sections of B16-OVA tumors harvested on day of sacrifice and stained with H&E and αCD8 in magenta. Scale bar equals 1cm for the tumor overview and 100 μm for the 20x magnification images. (J) Quantification of the number of $CD8^+$ infiltrating B16-OVA tumors based on tumor segmentation. One outlier in the IFs group was excluded, as tested by Grubb's outlier test with an alpha = 0.05.

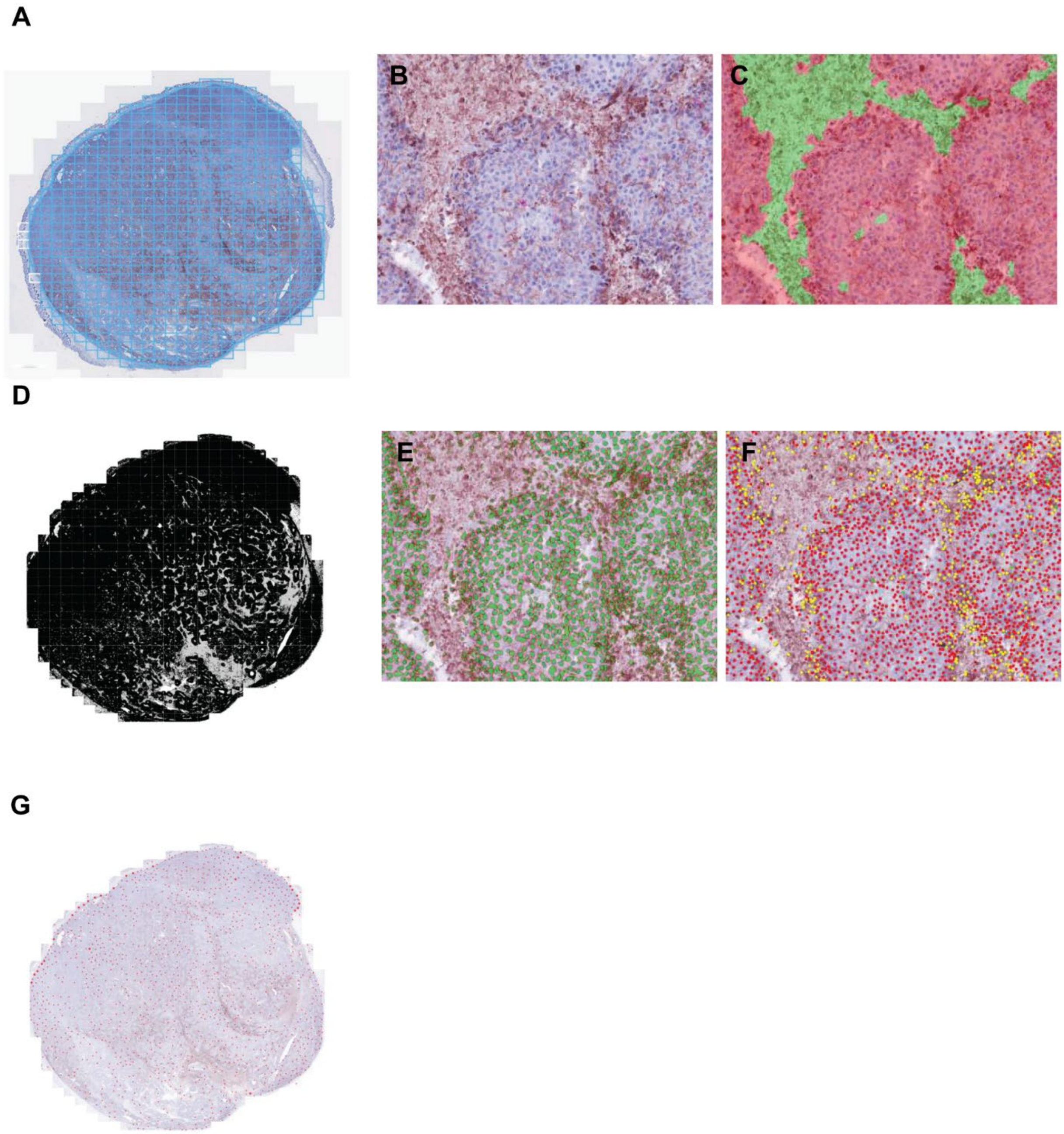


**Figure S10. Representative example of image analysis approach of T cell infiltration into B16-OVA tumors.** (A) B16-OVA tumors stained for CD8 were completely scanned at 20x magnification for multispectral imaging. (B) $CD8^+$ T cells (magenta), necrotic and pigmented regions (red-brownish) can be observed and could be spectrally unmixed. (C) A tissue segmentation algorithm was trained to separate tumor regions (red) from necrotic areas (green). (D) The tissue segmentation algorithm was applied to all images from all the samples to separate tumor regions (black) from necrotic areas (grey). (E) Cell segmentation was performed on the basis of hematoxylin staining. (F) A phenotyping algorithm was trained to recognize tumor cells (red dots), pigmented cells (yellow dots) and $CD8^+$ T cells (green dots). (G) The phenotyping algorithm was applied to all images from all the samples to quantify the density of $CD8^+$ T cells (red dots) in tumor region.

**Table S1**. **Normalized counts by RNAseq for all detected genes and of significantly different genes between DCs and IF-pMHC$^{(SIIT)}$/IL-2 after 8 hours and 22 hours stimulation of OT-I T cells.** (Separate excel file)
